# "Transcriptional Regulation of human NMNAT2: Insights from 3D Genome Sequencing and Bioinformatics"

**DOI:** 10.1101/2024.11.07.622375

**Authors:** Yu Chen Chang, Sen Yang, Minyoung Cho, Kwangsik Nho, José-Manuel Baizabal, Hui-Chen Lu

## Abstract

Nicotinamide mononucleotide adenylyl transferases 2 (NMNAT2) is a crucial nicotinamide adenine dinucleotide (NAD)-synthesizing enzyme essential for neuronal health. In the Religious Orders Study/Memory and Aging Project (ROSMAP), human brain levels of NMNAT2 mRNA positively correlated with cognitive capabilities in older adults. NMNAT2 mRNA abundance is significantly reduced following various insults or proteinopathies. To elucidate the transcriptional regulation of NMNAT2, we employed circular chromosome conformation capture followed by high-throughput sequencing (4C-seq) to identify potential NMNAT2 enhancer and silencer regions by determining genomic regions interacting with the NMNAT2 promoter in human SH-SY5Y cells. We discovered distinct NMNAT2 promoter interactomes in undifferentiated versus neuron-like SH-SY5Y cells. Utilizing bioinformatics analyses, we identified putative transcriptional factors and NMNAT2-associated genes. Notably, the mRNA levels of many of these genes showed a significant correlation with NMNAT2 mRNA levels in ∼400 single-nuclei RNA-seq datasets from ROSMAP. Additionally, using CRISPR-Cas9 strategies, we confirmed the requirement of two specific genomic regions within the interactomes and four transcription factors in regulating NMNAT2 transcription. In summary, our study identifies genomic loci containing NMNAT2 regulatory elements and predicts associated genes and transcription factors through computational analyses.

## Introduction

Cell type-specific transcriptomic analysis reveals the importance of transcriptional regulation in driving morphological and functional diversity in neurons [1]. Cells achieve differential gene expression by implementing complex gene regulatory networks [2]. The regulatory network often involves complex interplays between the three-dimensional (3D) organization of the target gene and its regulators in chromatin. The regulators include the genome’s cis-regulatory elements (CRE), trans-acting transcription factors (TFs), signaling molecules, and regulatory non-coding transcripts. CREs, such as enhancers and silencers, are located as close as a few base pairs or as far as millions of base pairs from the promoter region of target genes [3,4]. The 3D genomic organization enables long-distance CRE interaction with the promoter to regulate gene expression [5,6]. External insults or pathological conditions can change gene expression by altering chromatin structure or regulating TF abundance or activity. For example, widespread H3K27ac histone modifications, a robust marker for active CREs, were altered in the cortex of Alzheimer’s disease patients’ brains [7]. Additionally, axotomy results in large-scale changes in gene expression via injury-responsive TFs [8]. Several studies highlight the associations of transcriptional signatures with cognitive function [9], brain aging [10,11], and pathologic/clinical heterogeneity in neurodegenerative diseases [12–16].

Nicotinamide mononucleotide adenylyl transferases 2 (NMNAT2), a major nicotinamide adenine dinucleotide (NAD)-synthesizing enzyme in the mammalian brain [17], is critical for neuronal survival by maintaining axonal function [18–20]. Upon protein stress, NMNAT2 can also serve as a molecular chaperone to reduce protein aggregates [21]. Decreased NMNAT2 mRNA levels have been reported in the cortex of several neurodegenerative diseases, such as Alzheimer’s disease (AD), Parkinson’s disease (PD), and Huntington’s disease (HD) [21,11,22–24]. NMNAT2 transcription was also reduced by Wallerian-like axonal degeneration [25] and glaucoma [26,27], and over-expression of full-length human NMNAT2 exerts neuroprotective effects against glaucoma [28]. Deleting NMNAT2 in cortical neurons results in axonopathy and severe neurodegenerative-like phenotypes [18]. Previously, we found that NMNAT2 mRNA levels in the dorsolateral prefrontal cortex (DLPFC), a hub for cognitive and mood circuits, are positively correlated to cognitive capability before death in 541 deceased subjects collected during the Religious Orders Study and the Rush Memory and Aging Project (ROSMAP) [21]. Shigenoka et al. [29] showed that NMNAT2 mRNA is actively translated in axonal terminals, potentially serving as a constant supply of NMNAT2 protein. Furthermore, *NMNAT2*’s protein exhibits a very short half-life due to its targeted palmitoylation and is subject to the ubiquitin-proteosome pathway-mediated degradation [30]. Taken together, NMNAT2 mRNA transcription is a key step in regulating NMNAT2 protein abundance. A detailed understanding of NMNAT2 transcriptional regulation will facilitate the identification of risk factors for cognitive decline and the discovery of neuroprotection strategies.

To gain insights into the transcriptional regulation of NMNAT2, we leveraged the circular chromosomal conformation-sequencing (4C-seq) technique [31] to capture interactomes of the NMNAT2 promoter in the SH-SY5Y human neuroblastoma cell line in either an undifferentiated or differentiated neuron-like state. A CRISPR-Cas9 approach was employed to validate the selected interactomes and transcriptional factors. The genomic regions enrichment of annotations tool (GREAT) was employed to predict NMNAT2-associated genes with their putative cis-regulatory domains located in the genomic regions identified by 4C-seq as the interactomes with the NMNAT2 promoter. Next, we examined the relationship between NMNAT2 and predicted NMNAT2-associated genes using ROSMAP scRNA-seq data sets. The bioinformatic analysis also identified several putative transcriptional factors governing the expression of NMNAT2, including ATF4, ATF6, SOX11, and HSF1. Knocking down these transcriptional factors increased NMNAT2 mRNA levels and also altered the expression of EDEM3 and TPR, NMNAT2-associated genes identified from interactomes. Taken together, our study identified numerous putative enhancer/silencer genomic regions and transcriptional factors involved in human NMNAT2 gene transcription, as well as NMNAT2-associated genes.

## Material and Methods

### SH-SY5Y cell culture and differentiation

SH-SY5Y cells were cultured in a culture medium consisting of 45% eagle’s minimum essential medium (Sigma), 45% nutrient mixture F-12 Ham (Sigma), 10% heat-inactivated fetal bovine serum (FBS)(ATCC), 1% non-essential amino acids (Sigma), and 2 mM GlutaMax (Gibco). For differentiation, SH-SY5Y cells were seeded at a density of 10^4^ cells/cm in culture dishes coated with 0.05 mg/ml collagen (PureCol). Cells were treated with 10 μM all-trans-retinoic acid (RA) (Tocris) the next day. After 5 days of RA incubation, cells were washed with serum-free culture medium and subsequently treated with 50 ng/ml brain-derived neurotrophic factor (BDNF) (Alomone labs) in the serum-free culture medium for 7 days.

### Luciferase reporter assay

Two oligos with sequences corresponding to the two *in silico*-identified promoter regions (199 bp and 616bp) were synthesized with IDT gBlocks (IDT). These synthesized oligos were individually cloned into the pGL4.10 firefly plasmid (Promega) using HiFi DNA Assembly (NEB). pGL4.10 Firefly plasmid is a promoter-less vector for measuring the activity of promoter or enhancer sequences and contains Luc2, the second-generation version of firefly luciferase optimized for expression in mammalian systems. Cells were then transfected with Renilla and original or promoter-containing pGL4.10 Firefly plasmids in a 1:10 ratio using lipofectamine 3000 (Invitrogen). After 48 h, cells were then lysed and subjected to dual luciferase reporter assays (Promega) using the CLARIOstar plate reader (version 5.20 R3; BMG LABTECH). The firefly luminescence was normalized to the Renilla luminescence. To calculate relative fold changes, the normalized luminescence reads were divided by those from the empty vector.

### Chromosome conformation capture and 4C library preparation

We prepared the libraries according to a previously described protocol [32]. Briefly, 10 million cells were fixed with 2% formaldehyde (PFA) (Sigma) for 10 min at room temperature, quenched with cold 130 mM glycine (JT Baker), washed with cold Dulbecco’s phosphate-buffered saline (DPBS) twice to wash out the remaining PFA, and centrifuged at 500g 4°C for 5min and the supernatant discarded. The nuclear pellet was lysed in 1 ml lysis buffer (50 mM Tris-HCl pH 7.5, 0.5% IGEPAL CA-630, 1% Triton X-100, 150 mM NaCl, 5 mM EDTA, protease inhibitors) for 1h on ice. The lysed nuclear content was pelleted by centrifuge at 500g, 4°C, 5min, washed once with 1.2X restriction enzyme buffer, and the pellet resuspended in 500ul 1.2X restriction enzyme buffer. Subsequently, samples were treated with 0.3% SDS for 1h with gentle shaking at 900 rpm and 37°C, followed by 2.5% Triton X-100 for 1h with 900 rpm shaking at 37°C. Next, samples were treated with 600U of DpnII (NEB) and incubated overnight at 37°C with 900 rpm shaking. The following day, samples were subjected to heat inactivation at 65°C for 20 min to inactivate DpnII. Next, 50U T4 ligase (Roche, Cat.No. 10799009001) together with ligation buffer (66mM Tris-HCl, 5mM MgCl2, 5mm DTT, 1mM ATP, pH 7.5) was added to a final volume of 7ml, then, the samples were subjected to ligation at 16°C overnight. Cross-linking was reversed by adding 30 μl of protease K (10mg/ml; Sigma-Aldrich) at 65°C overnight and then treating with 30 μl of RNaseA (10mg/ml; Roche) for 45 min at 37°C. DNA was purified using Phenol-Chloroform-Isoamyl alcohol (25:24:1) (Sigma) with ethanol precipitation. The pellet was resuspended in 150 μl of 10 mM Tris-HCl (pH 7.5). Next, 50U of CviAII (NEB) in 10X restriction enzyme digestion buffer with a final volume of 500 μL was added for the second restriction enzyme digestion by incubating overnight at 25°C with 500 rpm shaking. Heat inactivation (65°C, 20 min) was performed to stop CviAII digestion. Samples were then ligated with 200U T4 ligase (Roche) in ml 10X ligation buffer diluted to a final volume of 14 ml at 16°C overnight. The ligated DNA was purified by ethanol precipitation, and the pellet resuspended in 150 μl of 10 mM Tris-HCl (pH7.5). QIAquick PCR purification kit (Qiagen Cat.No. 28104) was then used for final purification of the DNA. For preparing 4C-seq library, two PCR steps were performed. In the first step, 4 parallel PCR reactions with 200 ng of circularized DNA as the template (for each reaction) using Expand Long Template Polymerase mix (Roche) were performed with the primers as below: reading primer 5’-TACACGACGCTCTTCCGATCTCCCCTTAACGCTTGCAGC-3’, non-reading primer 5’-ACTGGAGTTCAGACGTGTGCTCTTCCGATCTCAAAAAGAGAGGCAACCCCTAG -3’. All PCR products were then pooled together and purified using AMPure XP beads (Beckman Coulter) according to the manufacturer’s Instructions. In the second step, PCR reactions with the previously purified PCR products as the template using Expand Long Template Polymerase mix (Roche) were performed with the primers as below: Universal primer 5’-AATGATACGGCGACCACCGAGATCTACA CTCTTTCCCTACACGACGCTCTTCCGATCT-3’, Indexed primer for biological replicate one 5’-CAAGCA GAAGACGGCATACGAGATCGTGATGTGACTGGAGTTCAGACGTGTGCT-3’, Indexed primer for biological replicate two 5’-CAAGCAGAAGACGGCATACGAGATACATCGGTGACTGGAGTTCAGACG TGTGCT-3’, Indexed primer for biological replicate three 5’-CAAGCAGAAGACGGCATACGAGATGCCTAAGTGACTGGAGTTCAGACGTGTGCT-3’. PCR products were then purified using QIAquick PCR purification kit (Qiagen). The DNA libraries were then sequenced using the Illumina NovaSeq 6000 SP platform.

### 4C-seq analysis

The single-end FASTQ raw data were analyzed using w4CSeq (github.com/WGLab/w4CSeq) and pipe4C pipelines (github.com/deLaatLab/pipe4C). Briefly, the raw reads were trimmed free of the viewpoint primer sequences, and then mapped to the reference genome hg19. In w4CSeq, the binomial model was used to call out statistically significant regions that pass p<0.05. For counting the number of ligated sites to define interchromosomal interactions, a window size of 500 was used. To define intrachromosomal interactions (shown as significant domains in Fig. 5), a window size of 100 with a background window size of 3,000 was used. FDR of 0.05 was used to remove most random chromosomal interactions. In pipe4C, the contact intensity was normalized to the fragment read counts (Sup. Fig. 4-1).

### Downstream analysis

Significant 4C interactomes were annotated to associated genes using the Genomic Regions Enrichment of Annotations Tool (GREAT). The basal plus extension option was used with the gene regulatory domain defined as 5 kb upstream and 1 kb downstream of the transcription start site with a maximum extension of 1,000kb.

Transcription factors of the 4C interactomes were predicted using WhichTF (bitbucket.org/bejerano/whichtf/src/master). The analysis was performed using the default setting in the pipeline without the need for user-defined criteria and cutoffs. The ranking order in the output of WhichTF is primarily used to evaluate prediction reliability, while the statistical P value is computed conditioned on the TF ranking. Transcription factors of the NMNAT2 promoter region (viewpoint) were predicted using the MEME Suite. Motif analysis was carried out using MEME version 5.5.3 (meme-suite.org/meme/tools/meme). Any number of repetitions was used for the distribution of motif sites in sequences. The discovered motifs were input in Tomtom version 5.5.3 (meme-suite.org/meme/tools/tomtom) using HOCOMOCOv11_core motif database and default parameters for transcription factor prediction. Transcription factor occurrences of HSF1, ATF4, ATF6, and SOX4 in region 1 and region 2 were predicted using FIMO version 5.5.5 (meme-suite.org/meme/tools/fimo) with default parameters. Motifs of HSF1, ATF4, ATF6, and SOX4 were retrieved from JASPAR 2022 motif database (jaspar2022.genereg.net).

### Cell type-specific gene-gene co-expression analysis using human single-nucleus RNA sequencing (snRNA-Seq) data

Cell type-specific gene-gene co-expression analysis between NMNAT2 and its associated genes was performed using single-nucleus RNA sequencing (snRNA-Seq) data previously generated from frozen samples of the dorsolateral prefrontal cortex (DLPFC) obtained from the brains of the Religious Order Study and the Memory Aging Project (ROS/MAP) participants (n=424) [33,34]. Pseudo-bulk expression levels were calculated. Briefly, individuals with fewer than 10 cells were excluded. A pseudo-bulk UMI count matrix was generated using the AggregateExpression function from the Seurat package [35]. Low-expression genes were filtered using the filterByExpr function from edgeR. TMM normalization (edgeR) was applied, followed by log2 transformation of CPM using the voom function from the limma [36,37]. Genes with log2CPM less than 2.0 were removed. Batch effects were corrected with ComBat and expression levels were normalized in quantile [38].

Pseudo-bulk expression level data in excitatory neurons and inhibitory neurons were used to examine the expression correlation between NMNAT2 and its associated genes. Pearson correlation coefficients were calculated using the cor.test function in R. Scatter plots for correlation visualization were generated using the ggplot2 package [39]. The false discovery rate (FDR) was computed to adjust for multiple comparisons across the correlation results.

### RNA-sequencing analysis

The total RNA of undifferentiated and differentiated SH-SY5Y cells was extracted following the manufacturer’s instruction using RNeasy Plus Universal Mini kit (Qiagen) and sent for RNA-sequencing with Illumina NovaSeq 6000 SP platform. Approximately 200 ng of total RNA was used for the library construction. Raw sequencing data was aligned to the human reference genome – hg19 and calculated the gene counts using STAR by default parameters. Uniquely mapped reads were retained for the subsequent analysis.

### External Data Resources

For identifying putative promoter regions of NMNAT2 and open chromatin area of 4C target regions, the ENCODE database was used to retrieve DNaseI data and ChIP-seq data of H3K27ac and H3K4me3 in brain-related samples. For DNaseI data, ENCFF877QYK, ENCFF714QMK, ENCFF175IJP, and ENCFF495MWO were used; for H3K27ac data, ENCFF506EFC, ENCFF036PHA, ENCFF500RRP, ENCFF534YU, ENCFF103SFW, ENCFF516KKW, ENCFF533OFX, ENCFF306NQW, and ENCFF297ASM were used; for H3K4me3 data, ENCFF586RDL, ENCFF433GYN, ENCFF478BHA, ENCFF080XAQ, ENCFF379BNK, ENCFF644BAH, ENCFF917YCH, ENCFF721PFP, ENCFF530API, ENCFF531LTO, and ENCFF565UAK were used. In addition, the conservation track (hg19.100way.phastCons.) from UCSC was used for comparison.

### piggyBac-based CRISPR/Cas9 gene deletion

CRISPR-Cas9 gene deletion for target regions and transcription factors was performed following the methods described in Schertzer et al. 2019. Briefly, gRNAs for deleting the target regions or transcription factors were designed and cloned into the pb_rtTA_Bsmb1 plasmid (Addgene) using Golden Gate Assembly (NEB). Cells were then co-transfected with the plasmid with gRNAs, the plasmid pb_tre_Cas9 (Addgene), and the plasmid piggyBacTransposase (Systems Bioscience) in a ratio of 8:2:1 using lipofectamine 3000 (Invitrogen). Two days after transfection, cells were treated with Hygromycin (400 μg/ml; Gibco) and G418 (700 μg/ml; Gibco) containing selection media. Following antibiotic selection, stable cell lines were maintained in media consisting of 45% Eagle’s minimum essential medium (Sigma), 45% nutrient mixture F-12 Ham (Sigma), 10% heat-inactivated FBS (ATCC), 1% non-essential amino acids (Sigma), 2 mM GlutaMax (Gibco), 400 μg/ml Hygromycin (Gibco) and 700 μg/ml G418 (Gibco) at 37°C and 5% CO_2_. For doxycycline induction, the stable cell lines were first seeded to 50% confluency and subsequently treated with 1 μg/ml of doxycycline (Millipore Sigma) for 4 days.

The gRNA sequences in this study are listed in Supplementary Table6. The primer sequences flanking the deleted genomic regions used for checking the efficiency of CRISPR-Cas9 deletion are listed in Supplementary Table6.

### RNA extraction and quantitative RT-PCR

The total RNA of SH-SY5Y or differentiated SH-SY5Y cells was extracted following the manufacturer’s instruction using the Quick-DNA/RNA MiniPrep Plus kit (ZYMO RESEARCH). 250 ng of total RNA was used for the reverse transcription of cDNA using TaqMan Reverse Transcription Reagents (applied biosystems). For comparison of gene expression, qRT-PCR was carried out using TaqMan Gene Expression Assays for NMNAT2 (Hs00322752_m1), EDEM3 (Hs00981767_m1), TPR (Hs00162918_m1), ODR4 (Hs00215258_m1), 18S (Hs99999901_s1), and GAPDH (Hs02758991_g1) in a 20 μl reaction consisted of 10 μl TaqMan Universal PCR Master Mix (applied biosystems), 1 μl TaqMan Gene Expression Assay, 100 ng of cDNA, and water, with the PCR program as follows: 40 cycles of 15 s at 95°C and 1 min at 60°C. For measuring gene expression of SOX11, HSF1, ATF4, and ATF6 relative to GAPDH as internal control (primer sequence in Supplementary Table6), qRT-PCR was conducted with duplicates in a 10ul reaction consisting of 5ul PowerUp SYBR Green Master Mix (A25742), 0.3ul 10µM forward and reverse primers, about 200 ng of cDNA, and water, following standard PCR program according to the manufactural instruction. All qRT-PCR were run in the QuantStudio™ 7 Pro Real-Time PCR Systems (Applied Biosystems).

### Quantification and statistical analysis

For each experiment, the number of biological replicates is indicated in the figure legend. Values are presented as mean ± SEM. Shapiro-Wilk normality test was used to check the normality of residuals. All data passed the normality test and thus two-tailed unpaired Student’s t-tests were performed to determine statistical significance.

## Results

### Identify the core promoter region for the human NMNAT2 gene

A promoter domain is a genomic area where RNA polymerase and transcription factors bind and initiate gene transcription [40]. Genomic sequences regulating gene transcription include not only the promoter upstream of the transcriptional start site but also enhancer/silencer elements that could be from a few thousand to millions of base pairs away from the gene target [41,3,4]. Transcription factors and co-activators or co-repressors bind to cis-regulatory elements (CRE) to form 3D structures to regulate gene expression. Recent advancements in chromosome conformation capture (3C) have allowed the identification of the interactive genomic loci that make contact with a specific locus of interest in nuclear space [42–44]. 4C-seq technology combines 3C principles with high-throughput sequencing to enable unbiased genome-wide screens for DNA contacts [45,46,44]. Thus, it allows the identification of 3D genomic interactions between enhancer/silencer and a single promoter region [47,48].

To employ 4C-seq to identify NMNAT2 enhancer/silencer regions, we started by exploring the putative *NMNAT2* promoter by examining chromatin marks upstream of the NMNAT2 transcriptional start site with DNase I hypersensitivity and H3K4me3 tracks listed in ENCODE database [49] as well as the UCSC conservation track (Fig. 1A). DNase I hypersensitivity tracks mark open chromatin regions, which are transcriptionally active. Histone H3 lysine 4 trimethylation (H3K4me3) marks active promoters. The UCSC conservation track (hg19.100way.phastCons.) indicates the measurements of evolutionary conservation among 100 vertebrate species. Promoter regions tend to be evolutionary conserved and thus we evaluated the conservation track near the NMNAT2 transcriptional start site. Combining the analysis of these features allowed us to narrow down the putative NMNAT2 core promoter region to 616 bp on chromosome 1 (Fig. 1A-B).

**Fig 1.**
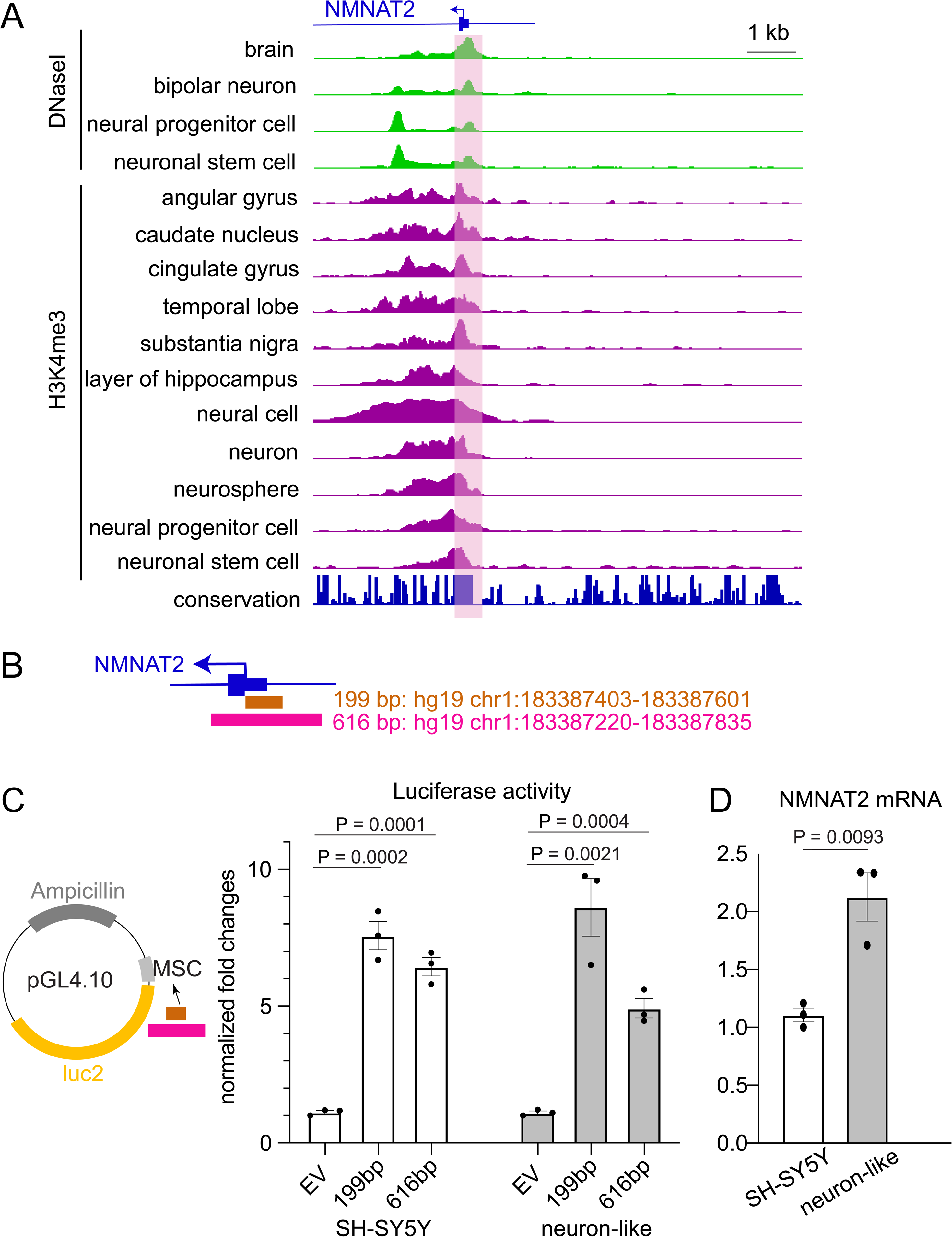
Identify the core promoter region of human NMNAT2. (**A**) Putative promoter regions of NMNAT2 – 199 bp and 616 bp – were identified through the chromatin marks, including DNaseI and H3K4me3, and the evolutionary conservation track (hg19.100way.phastCons.) retrieved from the ENCODE database. (**B**) Promoter regions of NMNAT2, 199 bp and 616 bp, and their corresponding chromosome locations (**C**) DNA fragments of 199 bp and 616 bp were cloned into the pGL 4.10 luciferase reporter vector using Hi-Fi Assembly. Luminescence signals from the promoter regions of NMNAT2 were normalized to the control, empty vector (EV) in SH-SY5Y (white) and differentiated SH-SY5Y (gray) cells. n = 3 independent batches of cells per group. (**D**) Summary of qRT-PCR for NMNAT2 mRNA levels in undifferentiated and neuron-like SH-SY5Y cells. n = 3 independent batches of cells per group.

To test the activity of the identified 616 bp putative NMNAT2 promoter region, we carried out dual luciferase promoter assays with SH-SY5Y cells. This human neuroblastoma cell line is derived from the sympathetic nervous system [50–52]. We detected significant luminescence when the 616 bp fragment was placed into the 5’ end of the promoter-less luciferase vector (Fig. 1C). Interestingly, within the 616 bp region, a sequence of 199 bp (chr1:183387403-183387601), immediately upstream of the transcriptional start site displays similar promoter activity (Fig. 1C).

SH-SY5Y cells can differentiate into neuron-like cells upon retinoic acid treatment for 5 days followed by 7 days of BDNF treatment [53]. NMNAT2 in human brain is mainly expressed in differentiated neurons, showing relatively low levels in neuronal progenitor cells [21,17]. To evaluate if NMNAT2 transcription is higher in neuron-like than undifferentiated SH-SY5Y cells, quantitative RT-PCR was conducted. We found a ∼2-fold increase of NMNAT2 mRNA levels in neuron-like compared to undifferentiated SH-SY5Y cells (Fig. 1D). Next, we tested the two putative promoter regions for their promoter activity with neuron-like SH-SY5Y cells. We found that the 199 bp sequence displays similar promoter functionality in neuron-like SH-SY5Y cells (Fig. 1C). However, the promoter activity of 616bp is significantly weaker in neuron-like SH-SY5Y cells compared to undifferentiated cells. These data suggest that in a neuron-like state, transcriptional suppressors of *NMNAT2* expression may bind outside of the 199 bp region but within the 616 bp promoter. Taken together, these experiments indicate that the 199 bp genomic region can serve as a robust functional NMNAT2 promoter.

### Identifying the interactomes of the NMNAT2 promoter in undifferentiated and neuron-like SH-SY5Y cells

The identification of the NMNAT2 promoter allows us to leverage 4C-seq technology [32,45,54] to identify the distal regulatory sequences interaction with the NMNAT2 promoter. To preserve as much as of the 199 bp sequence with optimal promoter activity as the viewpoint for comprehensive genome-wide interactome capture, DpnII and CviAII were selected as the restriction enzyme pair to prepare the 4C library, therefore rendering 186 bp uncut sequence out of 199 bp as the viewpoint (chr1:183387400-183387585) (Fig. 2A). We then carried out 4C-seq with both undifferentiated and neuron-like SH-SY5Y cells. Knowing that NMNAT2 transcription is significantly higher in a neuron-like state (Fig. 1D), we expected to identify different interactome sets in undifferentiated vs differentiated SH-SY5Y cells. Indeed, remarkably distinct NMNAT2 interactomes across the genome between these two states were identified by our 4C-seq (Fig. 2B & Table 1). Among 25 interactomes identified from undifferentiated SH-SY5Y cells, there are 9 intrachromosomal and 16 interchromosomal interactomes. For the 30 interactomes found with neuron-like SH-SY5Y cells, there are 17 intrachromosomal interactomes and 13 interchromosomal interactomes. Such large numbers of interchromosomal interactomes are consistent with previous studies [55,47]. Among these interactomes, only 2 intrachromosomal interactome regions partially overlapped between undifferentiated and neuron-like SH-SY5Y cells. The distinct profiles of interactomes suggest a differential combination of TFs and CREs regulating NMNAT2 transcription in un-differentiated and neuron-like SH-SY5Y cells.

**Fig 2.**
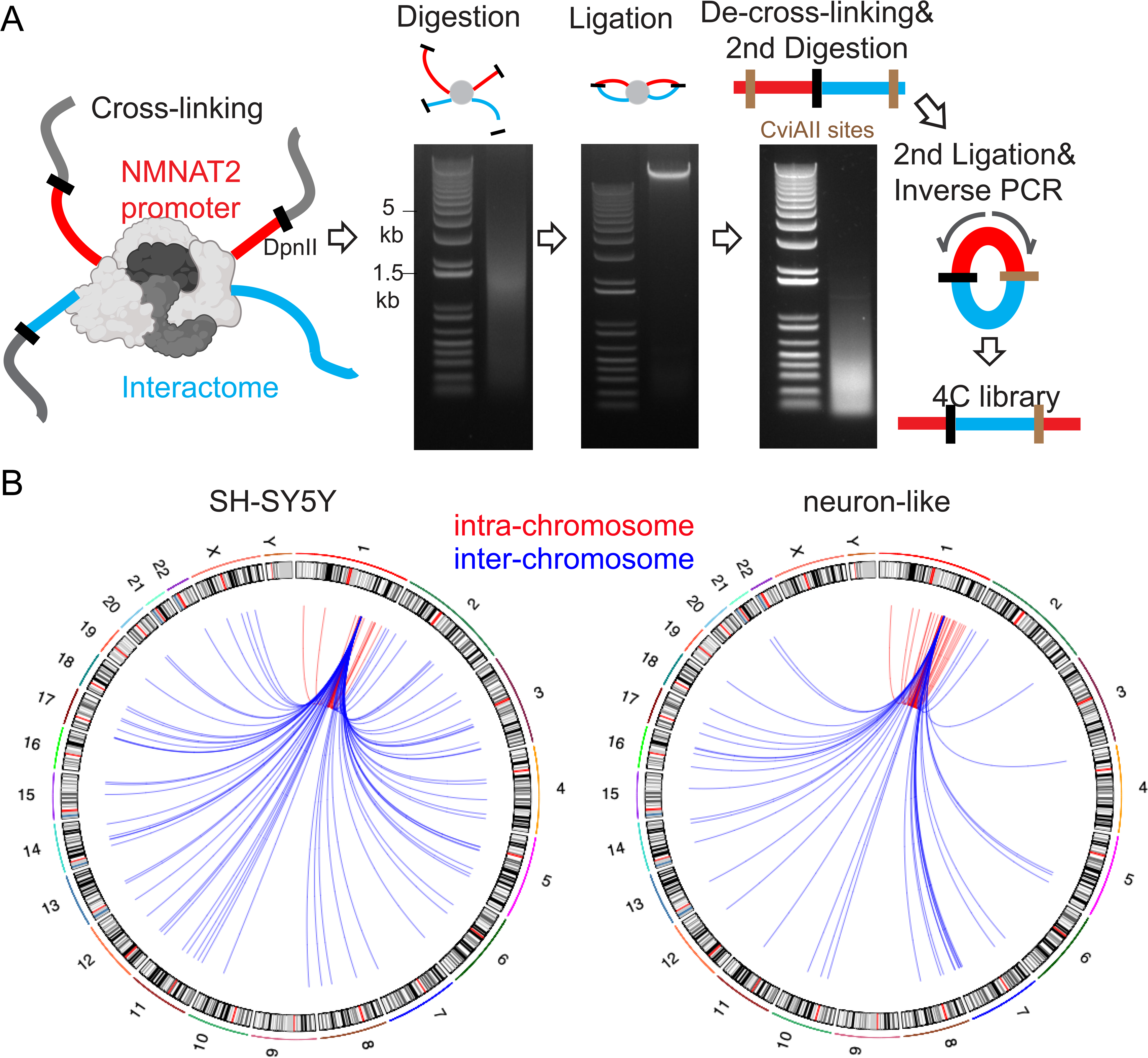
Identify 4C-sequencing interactomes using the NMNAT2 promoter as the viewpoint (bait). (**A**) Schematic of 4C-seq workflow. The NMNAT2 promoter was used as the viewpoint (bait) to capture genome-wide interactomes. Genomic regions that are spatially close in the cell nucleus were fixed by 2% formaldehyde via cross-linking. The DNA was first fragmented by DpnII and subsequently ligated by T4 ligase. De-cross-linking was applied to remove proteins. The purified DNA was then digested with CviAII and subsequently ligated by T4 ligase. An inverse PCR was carried out using the primers outward of the viewpoint to generate 4C library. The libraries were analyzed using next-generation sequencing Illumina NovaSeq 6000 platform. (**B**) Circos plots of intrachromosomal (red) and interchromosomal (blue) interactomes of the NMNAT2 promoter identified in SH-SY5Y and neuron-like SH-SY5Y cells using the w4Cseq pipeline analysis. Chromosomes are indicated around the circle. n = 3 independent batches of cells per group.

**Table 1.** 4C-seq intra-/inter-chromosomal interactomes. #: Overlapped intrachromosomal regions: intra11/intra12; intra15/intra16

| Cell Type | Region ID | Region Location |
| --- | --- | --- |
| SH-SY5Y | intra1 | chr1:81339897-81389305 |
|  | intra7 | chr1:178221095-178260036 |
|  | intra8 | chr1:179705026-179766034 |
|  | intra11# | chr1:184624695-184695665 |
|  | intra16# | chr1:186304238-186350135 |
|  | intra17 | chr1:186371783-186419706 |
|  | intra21 | chr1:216727072-216773062 |
|  | intra23 | chr1:226062672-226107828 |
|  | intra25 | chr1:245672841-245706016 |
|  | inter1 | chr2:9265434-9510208 |
|  | inter2 | chr2:66133777-66438510 |
|  | inter4 | chr3:118886471-119125972 |
|  | inter5 | chr3:183779352-183860069 |
|  | inter6 | chr4:119207589-119470595 |
|  | inter7 | chr4:159322784-159536468 |
|  | inter8 | chr5:137958668-138117199 |
|  | inter9 | chr7:30195666-30476677 |
|  | inter11 | chr7:133828443-134022198 |
|  | inter15 | chr8:128533899-128676811 |
|  | inter16 | chr9:8555947-8767604 |
|  | inter18 | chr11:34474688-34669429 |
|  | inter19 | chr12:19386382-19590374 |
|  | inter22 | chr15:90785383-90948054 |
|  | inter26 | chr17:29930733-30175550 |
|  | inter28 | chr18:25435-248817 |
| Neuron-like SH-SY5Y | intra2 | chr1:145659124-145690112 |
|  | intra3 | chr1:166248113-166289396 |
|  | intra4 | chr1:177087522-177155794 |
|  | intra5 | chr1:177254856-177296750 |
|  | intra6 | chr1:177559443-177597547 |
|  | intra9 | chr1:181063559-181107405 |
|  | intra10 | chr1:181953720-182030019 |
|  | intra12# | chr1:184639847-184730986 |
|  | intra13 | chr1:185097980-185158338 |
|  | intra14 | chr1:185243503-185306711 |
|  | intra15# | chr1:186272384-186375352 |
|  | intra18 | chr1:193094492-193147352 |
|  | intra19 | chr1:212138553-212199006 |
|  | intra20 | chr1:213122378-213161053 |
|  | intra22 | chr1:217017543-217059665 |
|  | intra24 | chr1:231532933-231588560 |
|  | intra26 | chr1:249134041-249178606 |
|  | inter3 | chr2:238229365-238435704 |
|  | inter10 | chr7:129502600-129650469 |
|  | inter12 | chr7:139334872-139626444 |
|  | inter13 | chr7:139815113-140045142 |
|  | inter14 | chr7:151785916-151993107 |
|  | inter17 | chr10:38988785-42734970 |
|  | inter20 | chr12:96474363-96643034 |
|  | inter21 | chr13:43868123-44094330 |
|  | inter23 | chr16:56442853-56656563 |
|  | inter24 | chr16:70710156-70902732 |
|  | inter25 | chr16:73328905-73519031 |
|  | inter27 | chr18:18272-180052 |
|  | inter29 | chr21:28216877-28445510 |

To test the function of the identified interactomes on NMNAT2 transcription, we employed the piggyBac-based CRISPR/Cas9 gene deletion method (Fig. 3A, B) to delete selected genomic loci and then examine their impact on NMNAT2 transcription. We selected the only two overlapping intrachromosomal interactome genomic loci identified in both undifferentiated and neuron-like SH-SY5Y cells (intra11/12 and intra15/16 in Table 1) for biological validation. These genomic regions cover the putative cis-regulatory domains of EDEM3, TPR, and ODR4 (Fig. 4A-C, Sup. Fig. 4-1). The identified significant interactome areas in these two genomic loci are large (∼50 kb) and difficult to delete with the piggyBac-based CRISPR-Cas9 method. Thus, we narrowed down the selection to include DNaseI tracks and the regulatory domains of EDEM3, TPR, and ODR4. Specifically, we selected the following regions to delete: (1) Region 1 is on chr1:184,664,369-184,669,585 (5218 bp) containing EDEM3’s regulatory domain with multiple DNase I tracks (Fig. 4A, Sup. Fig. 4-1A); (2) Region 2 is on chr1: 184,343,515-184,346, 086 (2573 bp) covering DNaseI tracks and the promoter region for TPR and ODR4 (Fig. 4B, Sup. Fig. 4-1B); (3) Region 3 is located on chr1:184662408-184664466 (2060bp) covering EDEM3 regulatory domain but lacking DNaseI tracks (Fig. 4C). Region 3 was designed to serve as a negative control for this deletion experiment. Genomic PCR experiments confirmed the success of the precise deletion of all three regions (Fig. 3B & Sup.-Fig. 3-1).

**Fig 3.**
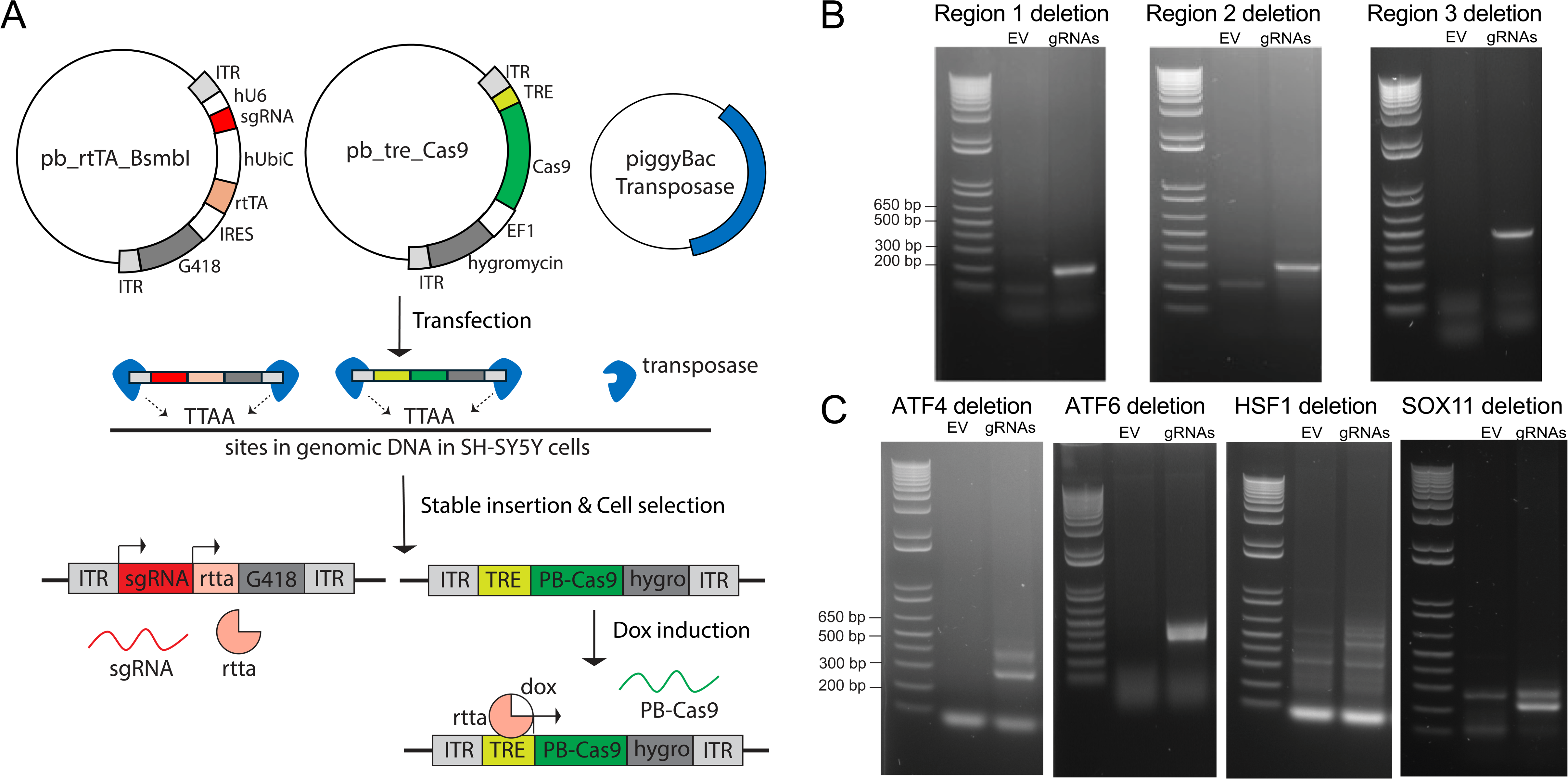
CRISPR-Cas9 deletion of the target regulatory regions and transcription factors. (**A**) Schematic of CRISPR-Cas9 deletion using piggyBac system. A Cas9-expressing piggyBac cargo vector (pb_tre_Cas9) is co-transfected with a gRNA-and rtTA-expressing piggyBac cargo vector (pb_rtTA_BsmbI) with a piggyBac transposase plasmid to generate a stable cell line through the hygromycin and G418 selection. ITR, piggyBac inverted terminal repeat. TRE, tetracycline responsive element. EF1, EF1a promoter. hU6, human U6 promoter. (**B**-**C**) Deletion check for the target regulatory regions (**B**) and transcription factors (**C**). PCR primers were designed flanking the deleted genomic sequences. The expected PCR product sizes were listed in Supplementary Table6 and Sup. Fig. 3-1. n = 3 independent batches of cells per group.

**Fig 4.**
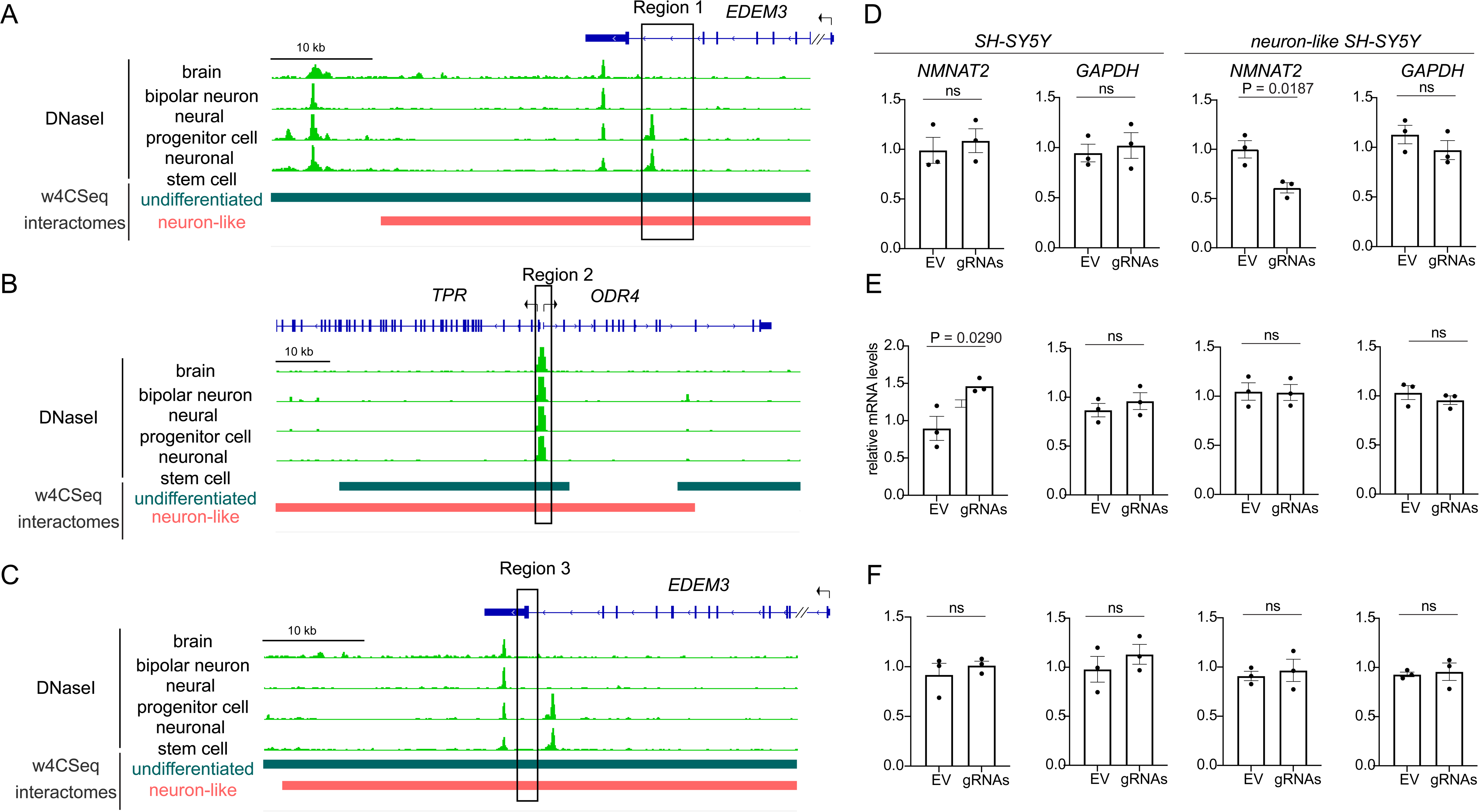
Genomic structure of selected interactomes and the impacts of their deletions on NMNAT transcriptions. (**A**) A cartoon illustrates the location of region1 (black box; chr1:184664369-184669585) relative to the regulatory domain of EDEM3 integrated with 4C interactomes output from the w4CSeq pipelines. (**B**) An illustration of the location of region2 (black box; chr1:186343515-186346086) relative to the regulatory domain of TPR and ODR4 integrated with 4C interactomes output from the w4CSeq pipelines. (**C**) An illustration showing the location of region3 (black box; chr1:184662408-184664466) relative to the regulatory domain of EDEM3 integrated with 4C interactomes output from the w4CSeq pipelines. (**D-F**) qPCR summary for the impact of regions 1-3 deletion on NMNAT2 mRNA levels in undifferentiated and neuron-like SH-SY5Y cells. (n = 3 independent batches of cells per group.

For SH-SY5Y cells with region 1 deleted, NMNAT2 mRNA was downregulated only in neuron-like but not in undifferentiated SH-SY5Y cells (Fig. 4D). In contrast, deleting region 2 resulted in a significant NMNAT2 upregulation in the undifferentiated but not the neuron-like SH-SY5Y cells (Fig. 4E). As expected, region 3 deletion exerted no impact on NMNAT2 expression in either state (Fig. 4F). These data suggest that regions 1 and 2 contain NMNAT2 CREs.

### Identify NMNAT2-associated genes with Genomic Regions Enrichment of Annotations Tool (GREAT) and examine their relationships with NMNAT2 mRNA levels in ROSMAP snRNA-Seq data sets

It has been widely observed that the coordinated activation or repression of a group of genes is localized either as a cluster or dispersed throughout the genome [56]. Such co-regulated expression requires the spatial recruitment of CREs and promoters of the co-regulated genes, as well as the partly shared TFs [57,58,56]. To determine the genes tentatively coregulated with NMNAT2, we searched for genes whose regulatory domains contain 4C-seq identified interactomes, using GREAT [59]. GREAT is a tool that first defines the regulatory domain of each gene across an entire genome, and subsequently computes the degree of association between the input genomic regions and the regulatory domain of a gene. Such GREAT analysis uncovered 39 NMNAT2-associated genes in undifferentiated SH-SY5Y cells and 31 NMNAT2-associated genes in neuron-like SH-SY5Y cells, with 8 overlapping genes (Fig. 5A, Tables 2-3).

**Table 2.** GREAT-annotated genes in SH-SY5Y cells.

| Gene Name | Chr. # | GREAT Rank | P-Value (-log10) | FDR Q-Value (-log10) | Observed Region Hits | Binom Region Set Coverage | Agora AD nominated genes |
| --- | --- | --- | --- | --- | --- | --- | --- |
| <u>EDEM3</u> | 1 | 1 | 242.67 | 238.4 | 90 | 23.5 | No |
| <u>C1orf21</u> | 1 | 2 | 235.62 | 231.65 | 90 | 23.5 | No |
| TOR1AIP2 | 1 | 3 | 231.56 | 227.77 | 72 | 18.8 | No |
| FAM163A | 1 | 4 | 209.15 | 205.49 | 72 | 18.8 | Yes |
| LEFTY1 | 1 | 5 | 110.19 | 106.62 | 32 | 8.36 | No |
| PYCR2 | 1 | 6 | 102.77 | 99.28 | 30 | 7.83 | No |
| TPR | 1 | 7 | 91.56 | 88.13 | 30 | 7.83 | Yes |
| <u>PRG4</u> | 1 | 8 | 57.38 | 54.01 | 27 | 7.05 | No |
| ENSG00000173213 | 18 | 9 | 49.9 | 46.58 | 19 | 4.96 | No |
| USP14 | 18 | 10 | 49.54 | 46.27 | 20 | 5.22 | No |
| ARHGAP31 | 3 | 11 | 46.49 | 43.27 | 19 | 4.96 | No |
| METTL14 | 4 | 12 | 45.06 | 41.88 | 21 | 5.48 | No |
| PRSS12 | 4 | 13 | 42.32 | 39.17 | 21 | 5.48 | No |
| OCLM | 1 | 14 | 32.42 | 29.29 | 12 | 3.13 | No |
| PDC | 1 | 15 | 25.92 | 22.83 | 12 | 3.13 | No |
| PLEKHA5 | 12 | 16 | 21.19 | 18.13 | 12 | 3.13 | No |
| ITGB1BP1 | 2 | 17 | 19.38 | 16.34 | 9 | 2.35 | No |
| SPRED2 | 2 | 18 | 18.34 | 15.33 | 12 | 3.13 | No |
| AEBP2 | 12 | 19 | 18.21 | 15.22 | 12 | 3.13 | No |
| TMEM39A | 3 | 20 | 17.72 | 14.75 | 8 | 2.09 | No |
| ASAP2 | 2 | 21 | 16.97 | 14.02 | 9 | 2.35 | No |
| COPRS | 17 | 22 | 16.13 | 13.21 | 9 | 2.35 | No |
| RAB11FIP4 | 17 | 23 | 15.96 | 13.05 | 9 | 2.35 | No |
| MEIS1 | 2 | 24 | 15.84 | 12.95 | 12 | 3.13 | No |
| B4GALT4 | 3 | 25 | 13.76 | 10.89 | 6 | 1.57 | No |
| SMYD3 | 1 | 26 | 11.49 | 8.64 | 8 | 2.09 | No |
| KIF26B | 1 | 27 | 11.1 | 8.26 | 8 | 2.09 | No |
| ZNRF2 | 7 | 28 | 10.93 | 8.11 | 6 | 1.57 | No |
| RXFP1 | 4 | 29 | 10.18 | 7.38 | 6 | 1.57 | No |
| HTR3E | 3 | 30 | 9.36 | 6.56 | 4 | 1.04 | No |
| ODR4 | 1 | 31 | 8.22 | 5.45 | 3 | 0.78 | No |
| NOD1 | 7 | 32 | 7.53 | 4.77 | 4 | 1.04 | No |
| EIF2B5 | 3 | 33 | 7.33 | 4.58 | 3 | 0.78 | No |
| TMEM144 | 4 | 34 | 6.78 | 4.04 | 4 | 1.04 | No |
| GABARAPL3 | 15 | 35 | 6.58 | 3.85 | 3 | 0.78 | No |
| ELF5 | 11 | 36 | 5.59 | 2.88 | 3 | 0.78 | No |
| EHF | 11 | 37 | 4.63 | 1.93 | 3 | 0.78 | No |
| HTR3C | 3 | 38 | 4.47 | 1.78 | 2 | 0.52 | No |
| NGRN | 15 | 39 | 4.08 | 1.4 | 2 | 0.52 | No |

**Table 3.**
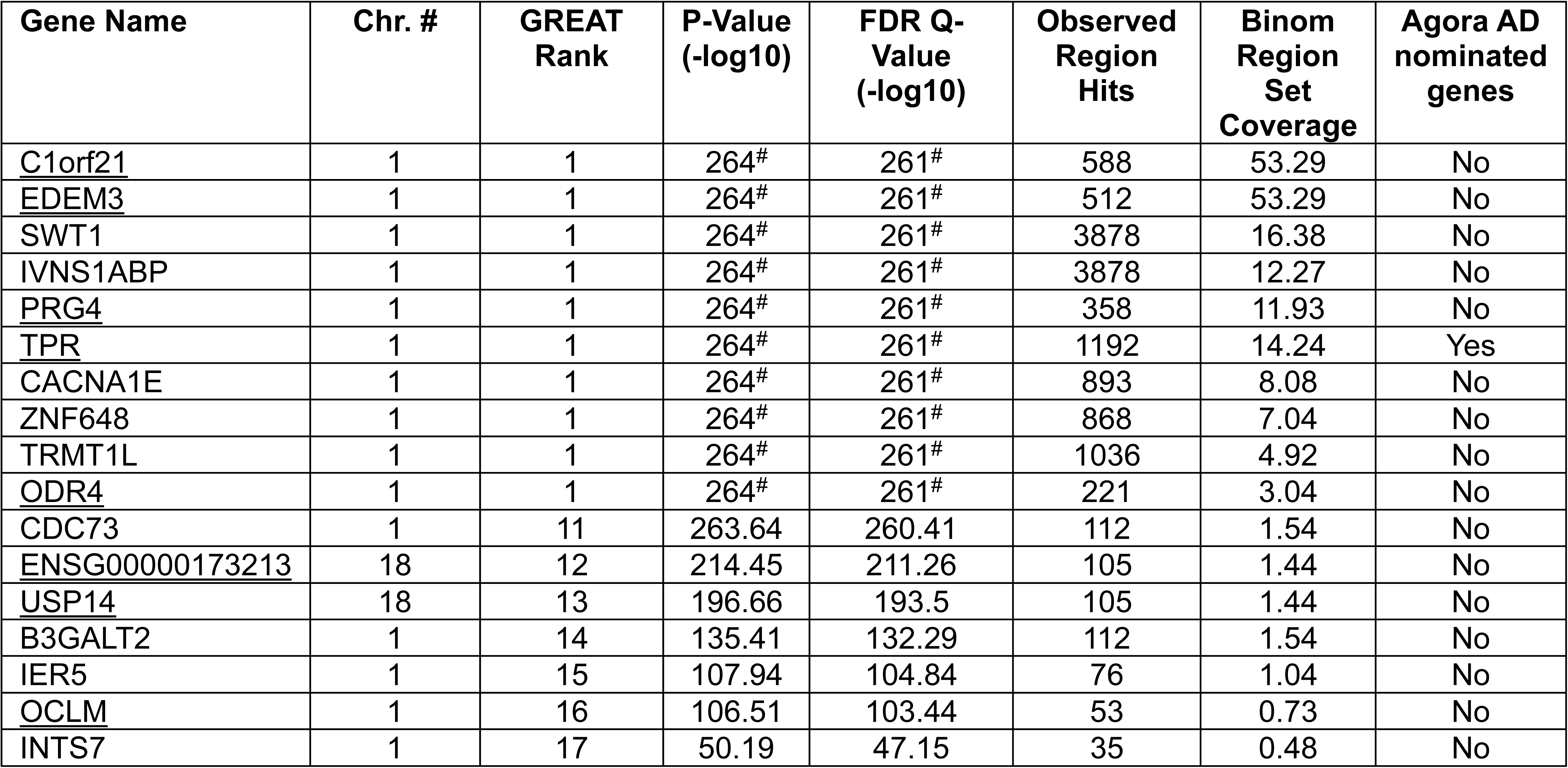
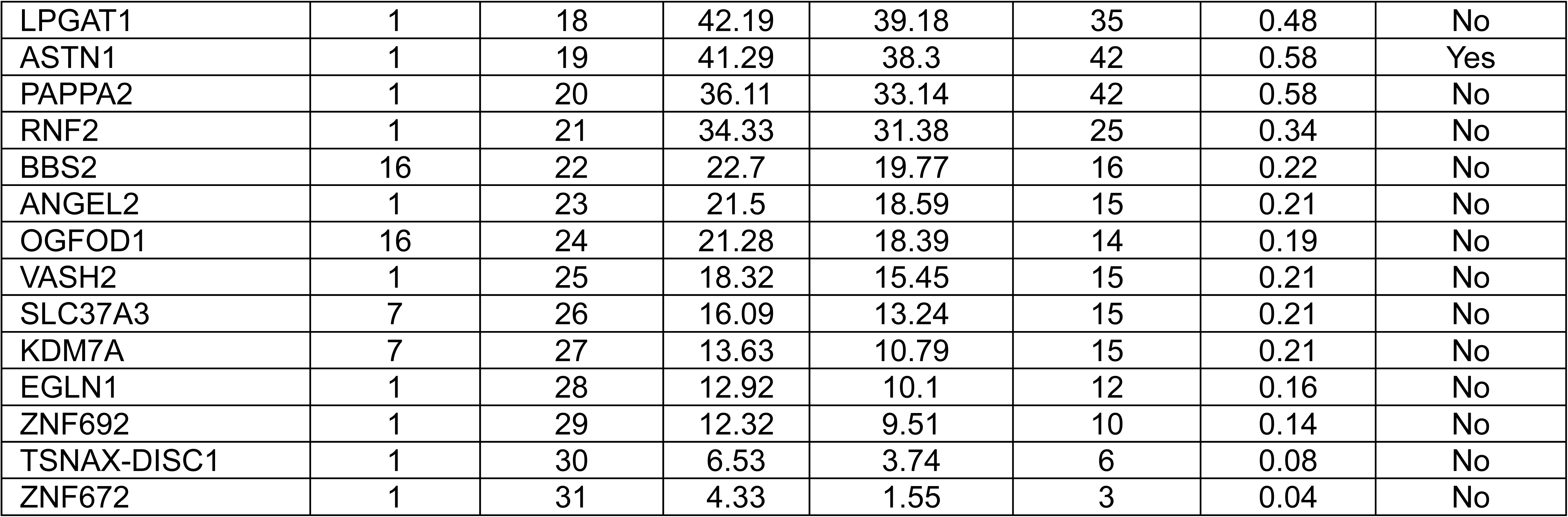
GREAT-annotated genes in Neuron-like SH-SY5Y cells. #: P-value and FDR are highly significant and close to 0, which is smaller than the precision of GREAT calculation.

**Fig 5.**
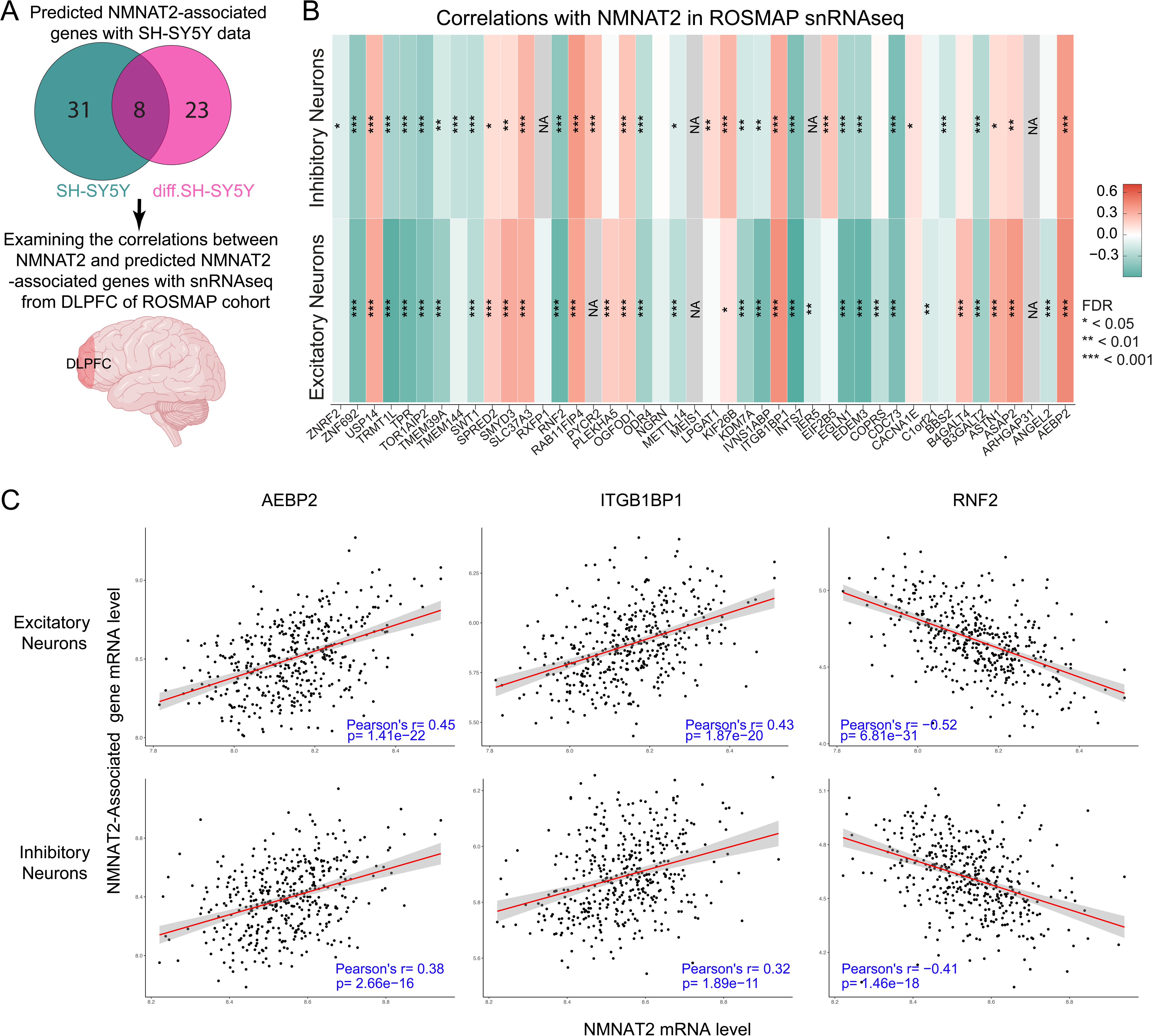
Coexpression of NMNAT2 and the predicted associated genes in aged human subjects. (**A**) Venn diagram shows the number of predicted NMNAT2 genes from informative analyses with interactomes. The relationships between NMNAT2 and these putative NMNAT2-associated genes were examined with ROSMAP snRNA-Seq data. (**B**) Heatmap shows the degree of Pearson correlation between NMNAT2 and NMNAT2-associated genes. Colors represent the correlation R values. Significance levels were denoted as follows: * FDR < 0.05, ** FDR < 0.01, *** FDR < 0.001. (**C**) Scatter plots for the distributions of AEBP2, ITGB1BP1, and RNF2 relative to NMNAT2 by their mRNA levels. Each dot represents data from one human subject.

Previously, we found that NMNAT2 mRNA levels in the dorsolateral prefrontal cortex (DLPFC), a hub for cognitive and mood circuits, of 541 deceased ROSMAP subjects were positively correlated to cognitive capability before death [21]. Here we took advantage of the recently released single nuclei (sn)RNA-Seq data from the DLPFC of 424 ROSMAP human subjects [33,34] to analyze the relationship between NMNAT2 mRNA levels and GREAT-predicted NMNAT2-associated genes. We focused on excitatory and inhibitory neurons because NMNAT2 expression is relatively abundant in neurons but low in astrocytes, microglia, immune, and vascular cell types. Pseudo-bulk expression level data in excitatory and inhibitory neurons were used to calculate the correlation coefficients for the expression levels of NMNAT2 and individual NMNAT2-associated genes (Table 4, Fig. 5B and Sup. Fig. 5-1).

**Table 4.**
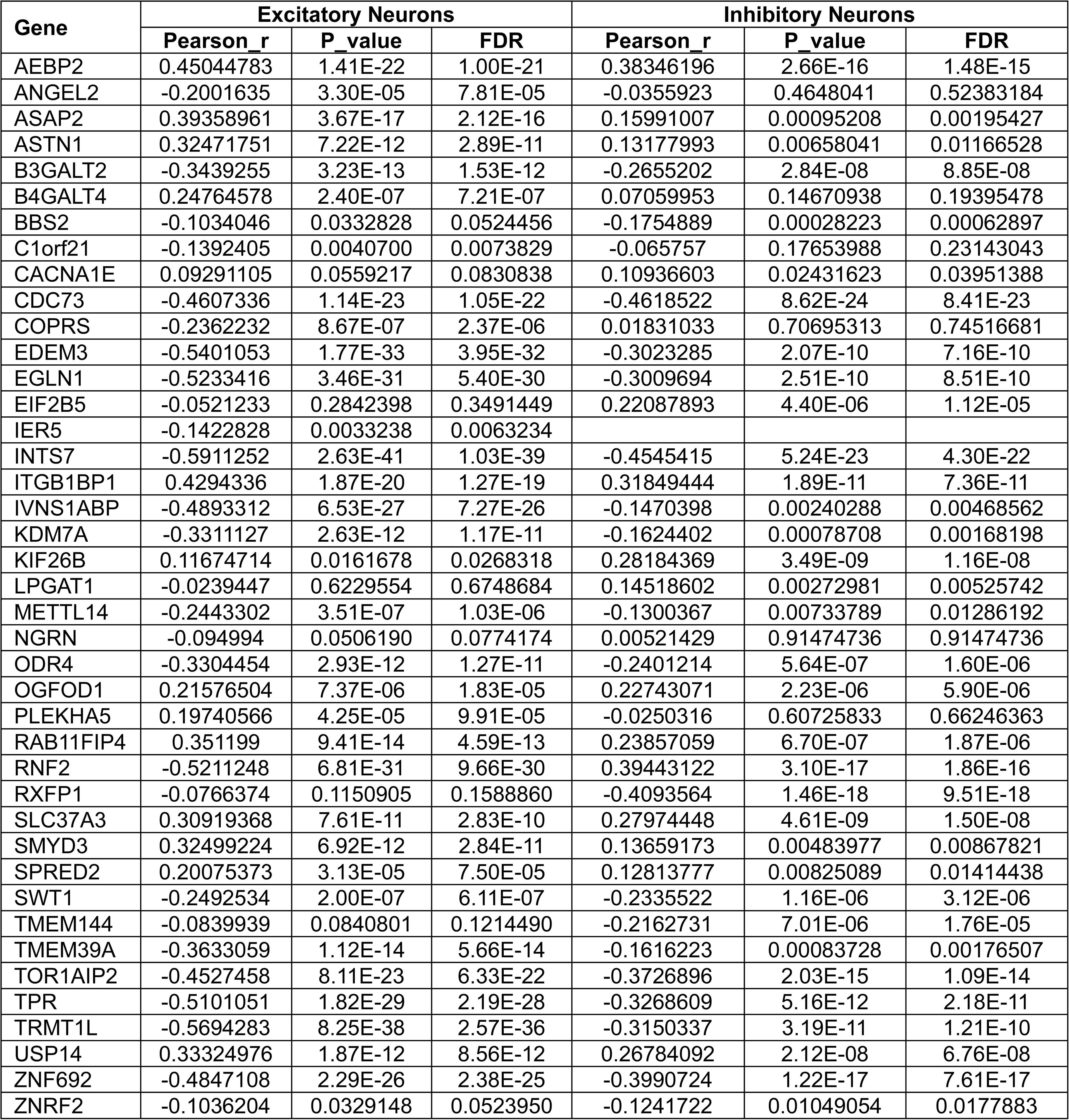
Co-expressed relationships between NMNAT2 and its associated genes in ROSMAP sn-RNAseq data.

The pseudo-bulk expression levels of 44 out of 62 NMNAT2-associated genes identified above (Tables 2-3) in ROS-MAP snRNA seq data sets are above the expression threshold (see Methods). Notably, we found the expression levels of 75% (33 out of 44) and 77% (34 out of 44) of NMNAT2-associated genes are significantly correlated with NMNAT2 mRNA levels in excitatory and inhibitory neurons, respectively (Fig. 5B). For example, the expression levels of AEBP2 and ITGB1BP1 are significantly positively correlated to NMNAT2 mRNA levels, while RNF2 mRNA levels are significantly negatively correlated to NMNAT2 mRNA levels (Fig. 5C). The directionality of correlations in excitatory and inhibitory neurons are mostly similar (Table 4 and Sup. Fig. 5-1). The finding of the correlated expressions between NMNAT2 with such a significant number of NMNAT2-associated genes in aged human subjects provides strong support for the presence of the 3D organization formed between NMNAT2 promoter regions with 4C-seq-identified interactomes. Furthermore, NMNAT2 abundance is co-regulated with many genes in neurons.

### Identify the putative transcriptional factors regulating NMNAT2 transcription via WhichTF and MEME Suite

Transcriptional factors (TFs) contain DNA binding domains and bind to specific DNA motifs in the promoter and/or CRE to modulate gene transcription [5,60]. The significant correlations between NMNAT2 transcription and most GREAT-predicted NMNAT2-associated genes suggest that TFs form complexes with both the NMNAT2 promoter and interactome where the regulatory regions for NMNAT2 associated genes are located and thus co-regulate the transcription of NMNAT2 and its associated genes. To identify putative TFs engaged in regulating NMNAT2 transcription, TF prediction analyses were conducted by searching their binding motifs in the genomic regions of the NMNAT2 promoter and the 4C-seq captured interactomes. Specifically, MEME Suite [61] was used to identify putative TFs directly bound to NMNAT2 promoter regions (Fig. 6A), while WhichTF pipeline [62] was applied to predict TFs interacting with NMNAT2 promoter regions through their binding with the 4C-interactomes (e.g. Fig. 6B).

**Fig 6.**
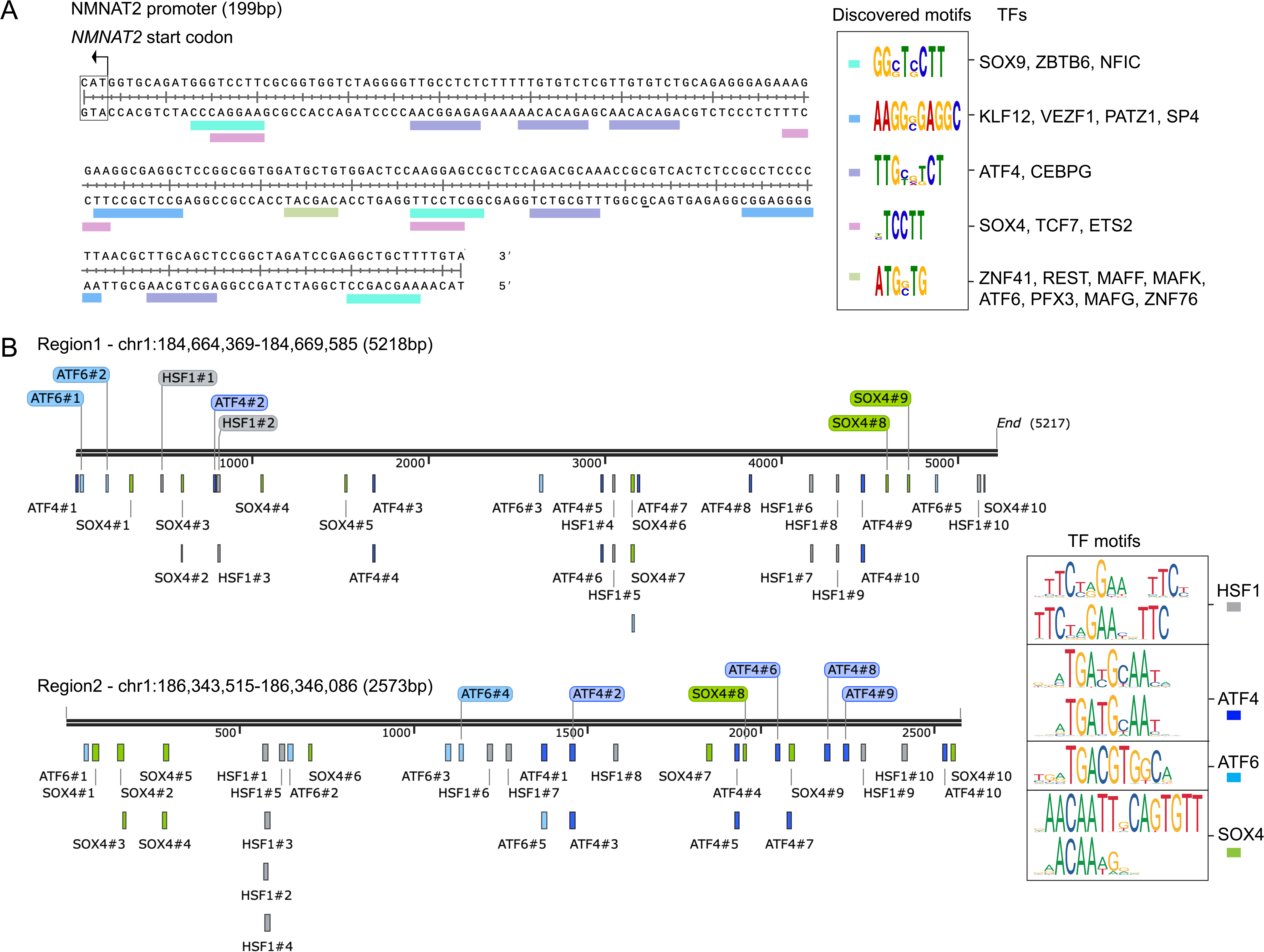
Transcription factor prediction of the NMNAT2 promoter and transcription factor occurrences of the region1 and region2. (**A**) Top20 transcription factors and their motif sequences in the NMNAT2 promoter. (**B**) Motif occurrences of HSF1, ATF4, ATF6, and SOX4 in region1 and region2.

MEME Suite identified 50 TFs that bind to the NMNAT2 promoter region. Based on the RNAseq data acquired from undifferentiated and neuron-like SH-SY5Y cells (Sup. Table 1), 20 out of 50 TFs are significantly expressed. Ranked by p-values, these 20 TFs include: ZNF41, REST, MAFF, SOX9, ZBTB6, KLF12, SOX4, ATF4, TCF7, CEBPG, MAFK, VEZF1, NFIC, ETS2, ATF6, RFX3, PATZ1, MAFG, ZNF76, and SP4 (Fig. 6A; Sup. Table 2). WhichTF integrates functional annotations from GREAT and the prediction of conserved human TF binding sites from PRISM to compute a stratified enrichment statistic, namely WhichTF score, to predict functionally dominant TFs interacting with our 4C-interactomes [62]. After excluding the predicted TFs with undetectable or low-level expressions (log10CPM < 1), we identified 247 functionally dominant TFs common to both undifferentiated and neuron-like SY5Y cells, 14 unique TFs in undifferentiated cells, and 11 unique TFs in neuron-like SY5Y cells (Sup. Table 3).

### ATF4/6, HSF1, and SOX11 regulate the transcription of NMNAT2 and its-associated genes

To validate our *in-silico* predictions, we selected ATF4, ATF6, HSF1, and SOX11, predicted TFs, to experimentally validate their roles in NMNAT2 transcription. We selected ATF4 and ATF6 because of their binding motifs on the NMNAT2 promoter. HSF1 was one of the predicted TFs from 4C-interactomes and interacted with TPR [63], a NMNAT2-associated gene tested above (Tables 2-3). SOX11 was selected for its high rank in WhichTF outputs and its conserved motif with SOX4 [64], which was predicted to bind to the NMNAT2 promoter. ATF4, ATF6, and HSF1 are more abundant in un-differentiated than neuron-like SH-SY5Y cells (Sup. Table 3). We employed the CRISPR-Cas9 method to knock down these TFs individually in SH-SY5Y cells (Fig. 3C) and then conducted qPCR to quantify NMNAT2 mRNA (Sup. Fig. 3-2). We found that ∼45% ATF4, ∼20% ATF6, ∼30% SOX11, or ∼40% HSF1 reduction led to significant upregulation of NMNAT2 expression in undifferentiated SH-SY5Y cells (Fig. 7; Sup. Fig. 3-2). In the neuron-like SH-SY5Y cells, ∼30% ATF4 or ∼25% HSF1 reduction also modestly, but significantly increased NMNAT2 expression, whereas ∼25% ATF6 or ∼30% SOX11 reduction does not affect NMNAT2 mRNA levels (Sup. Fig. 7-1).

**Fig 7.**
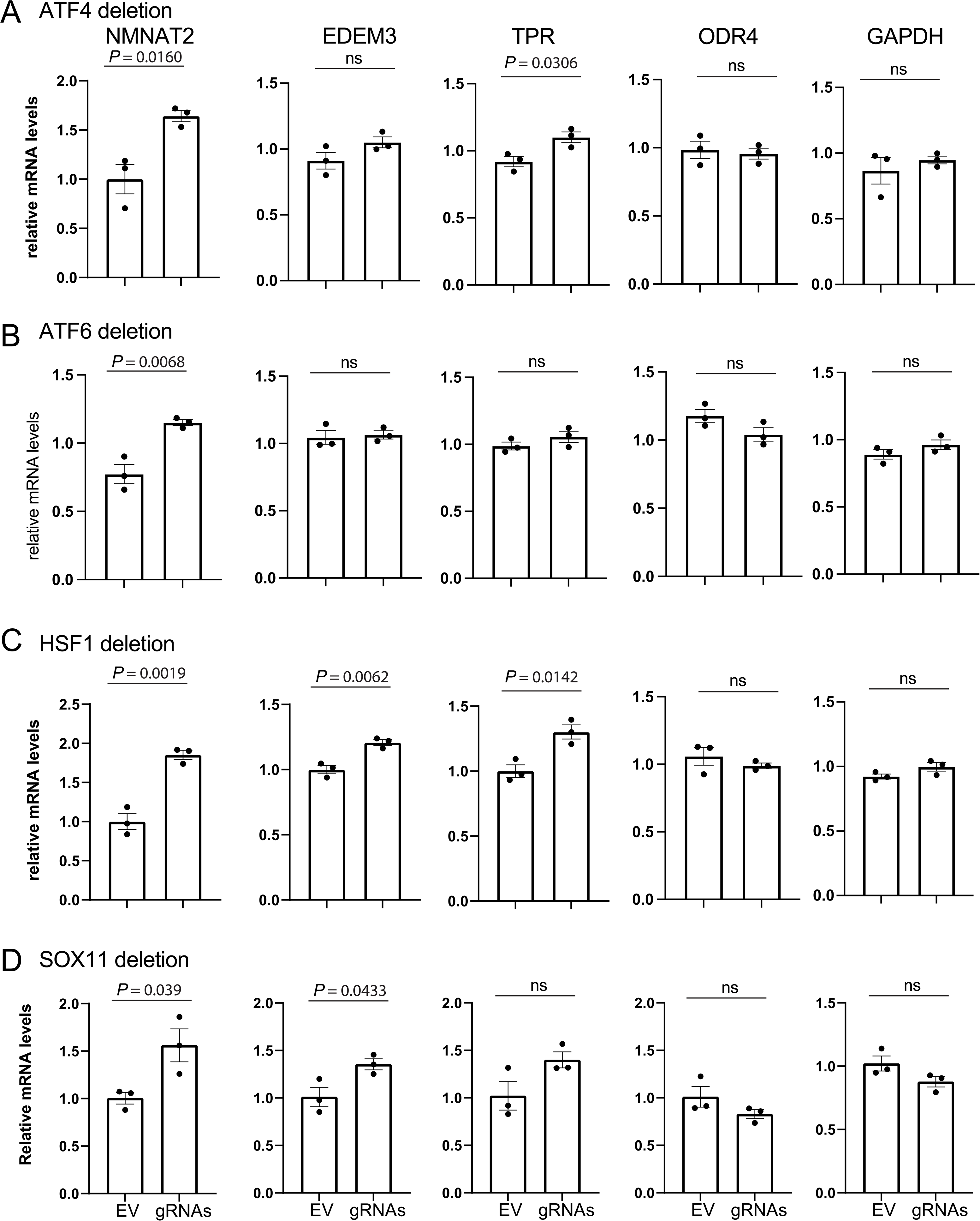
Impact of ATF4 (A), ATF6 (B), HSF1 (C), and SOX4 (D) knockdown on the expression levels of NMNAT2 and EDEM3, TPR and ODR4, three selected NMNAT2-associated genes in undifferentiated SH-SY5Y cells. n = 3 independent batches of cells per group.

Next, we performed motif occurrence analysis (FIMO) [65] with genomic region 1 and region 2 examined above, we found that ATF4, ATF6, and HSF1 have significantly enriched motif occurrences in these regions (Fig. 6B, Sup Tables 4-5). SOX11 motif sequence is lacking in the current database. Since SOX11 shares high similarity in the motif sequences with SOX4 in mice [64], we thus performed FIMO analysis for SOX4 and found that SOX4 also has significantly enriched motif occurrences in region1 and region2. The outcome of FIMO analyses raises the possibility that ATF4, ATF6, HSF1, and SOX11 bind onto region1 and region2. These regions cover the regulatory domains of EDEM3, TPR, and ODR4. Thus, we evaluated the mRNA levels of EDEM3, TPR, and ODR4 in ATF4/6, SOX11, HSF1 knockdown SH-SY5Y cells. In undifferentiated SH-SY5Y cells, we found that ATF4 knockdown increased expression of TPR (Fig. 7A). HSF1 knockdown increased EDEM3 and TPR mRNAs (Fig. 7C). SOX11 knockdown increased EDEM3 expression (Fig. 7D). Deleting ATF6, however, did not change EDEM3, TPR, or ODR4 expression (Fig. 7B). In neuron-like SH-SY5Y cells, both ATF4 and HSF1 knockdown increased EDEM3 with no change in TPR or ODR4 (Sup. Fig. 7-1 A-B). These data suggest that both ATF4 and HSF1 repress the expression of NMNAT2 and EDEM3. In undifferentiated SH-SY5Y cells, ATF4 and HSF1 also regulate TPR expression.

**Table 5.** Conditions of altered NMNAT2 expression.

|  | Induction/reduction in NMNAT2 | Condition/treatment | Samples/subjects | Reference |
| --- | --- | --- | --- | --- |
| **Humans** | 25-50% mRNA reduction | Diabetic retinopathy (DR) | Peripheral blood mononuclear cells from DR patients (n=52) and healthy controls (n=27) | [104] |
|  | 10-55% mRNA reduction | Parkinson's disease | Substantia nigra in microarray datasets from (GSE7621:PD n=16 and controls n=9; GSE20168:PD n=14 and controls n=15) | [105] |
|  | 60-80% mRNA reduction | Depletion of mitochondrial deoxyguanosine kinase (DGUOK) | Lung cancer cell lines (H1650; H1299) (n=3 for treatment and control groups) | [106] |
|  | 25-50% mRNA reduction | iPSC-derived neurons treated with niclosamide or static (STAT3 inhibitors) | iPSC-derived neurons (n=3 for treatment and control groups) | [107] |
|  | mRNA reduction | Breast cancer | mRNA and miRNA data of breast cancer from TCGA cancer genome database | [108] |
|  | 65-85% reduction in protein levels | Glaucoma | Ganglion cell complex (GCC) from glaucoma eyes (n=7) and control eyes (n=11) | [26] |
|  | 2 fold mRNA induction | Ovarian cancer samples | Ovarian cancer tissues (n=426) and ovarian normal tissues (n=88) | [109] |
| | 1.7 fold mRNA induction | iPSC-derived neurons treated with 100ng/ml IFN-γ for 48 h | iPSC-derived neurons (n=3 for treatment and control groups) | [107] |
|  | NMNAT2 positive occurrence (82.1%) in tumor tissues were higher than adjacent normal tissues (35.7%) [protein levels] | Colorectal cancer (CRC) | Tumor tissues from colorectal cancer (CRC) patients (n=95) | [71] |
| **Animals** | 40% mRNA reduction | Glaucoma mouse model induced by silicone oil-induced ocular hypertension (SOHU) | Glaucomatous retinal ganglion cells (RGCs) and normal RGCs (n=3 for each group) | [27] |
|  | 75% mRNA reduction | Mouse aged oocytes | Young/ aged mouse oocytes (n=50 for each group) | [110] |
|  | 50%-70% mRNA reduction | sALS mouse model – wobbler mice | Mouse spinal cord tissues (n=3) | [111] |
|  | 50% mRNA reduction | Depletion of SIRT1 | Differentiated 3T3-L1 cells (n=4) | [112] |
|  | mRNA reduction | Wallerian-like axon degeneration model induced by tri-ortho-cresyl phosphate (TOCP) | Chicken embryonic dorsal root ganglia (DRG) | [25] |
|  | 60% reduction in protein levels | Parkinson's disease rat model induced by 6-OHDA | Brain tissues of 6-OHDA-lesioned rats (n=8) | [105] |
|  | 1.5 fold mRNA induction | Adipogenesis of mouse 3T3-L1 cells | Differentiated 3T3-L1 cells (n=3) | [113] |
|  | 3 fold mRNA induction | Young and aged mouse islets | Young and aged mouse islets (n=3-4) | [114] |
| | 4-15 fold mRNA induction | Depletion of glutamate-oxaloacetate transaminase 1 (GOT1) in mouse clonal β-cell MIN6-K8 cells | Mouse clonal β-cell MIN6-K8 cells (n=3) | [114] |
| | 3 fold induction in protein levels | Neonatal rat cardiomyocytes treated with EGCG (major component in green tea) 50μM for 48h | Neonatal rat cardiomyocytes (n=5 for treatment and control groups) | [115] |
| | 2 fold induction in protein levels | Mouse primary astrocytes under the oxygen-glucose deprivation/reoxygenation (OGD/R) insult treated with 5μM ginsenoside Rb1 (major component in ginseng) | Mouse primary astrocytes (n=3) | [116] |

## Discussion

NMNAT2 is the major NAD-synthesizing enzyme in CNS neurons [66]. Low NMNAT2 increases neuronal susceptibility to neurodegenerative insults [67,68]. Many studies show that various pathological conditions can result in reduced NMNAT2 mRNA levels, such as nerve injury, proteinopathy, and glaucoma (Table 5), while NMNAT2 overexpression is neuroprotective against these insults [69,27,28]. Thus, NMNAT2 has been considered as a valuable drug target [70,71,28]. To elucidate the transcriptional regulation on NMNAT2 expression, we conducted a 4C-seq experiment to unbiasedly identify genomic regions containing NMNAT2 regulatory elements. Upon differentiation into the neuron-like state, SH-SY5Y cells express significantly higher NMNAT2 mRNA than at an undifferentiated state making them a useful system to study NMNAT2 transcriptional regulation. Interestingly, distinctive sets of interactomes were identified for the two different states, with more intrachromosomal interactomes found in neuron-like than undifferentiated SH-SY5Y cells. These findings suggest a change in 3D interactions among TFs and CREs to enhance neuronal NMNAT2 transcription upon differentiation. GREAT analyses with interactomes in both the undifferentiated and neuron-like states identify 62 NMNAT2-associated genes. Excitingly, we found significant relationships in expression levels between NMNAT2 and >75% of the GREAT-identified NMNAT2-associated genes with snRNAseq data from 424 ROSMAP human brains. Using CRISPR-Cas9 to selectively delete/knockdown selected genomic regions and ATF4, ATF6, SOX11, and HSF1, we confirmed their modulatory roles on NMNAT2 transcriptions.

### The roles of NMNAT2-associated genes in neuronal physiology

In this study, we identified several NMNAT2-associated genes in undifferentiated and neuron-like SH-SY5Y cells. Among these, we found that ∼70% of them display significant relationships with NMNAT2 expression levels in the neurons from the aged prefrontal cortex (Fig. 5). A few NMNAT2-associated genes implicated in neuronal function in the literature are worth mentioning here. Endoplasmic reticulum-associated protein degradation (ERAD)-enhancing α-mannosidase-like protein (EDEM3) and translocated promoter region protein (TPR) have been reported to be neuroprotective. EDEM3 is found to recognize and direct misfolded proteins to the ERAD process and subsequently protect cells from ER stress [72]. It is shown that upon aging, the activity of the ERAD process is significantly reduced and EDEM3 upregulation mitigates ER proteinopathy [73]. TPR is one of the crucial components of the nuclear pore complex (NPC) [74] and has been linked to neurodegenerative diseases, such as amyotrophic lateral sclerosis (ALS) and Huntington’s disease (HD). For instance, TPR was significantly down-regulated in human-induced pluripotent stem cell-derived motor neurons (iPSNs) harboring the C9ORF72 mutation, the most common cause of familial ALS [75]. TPR interacts with the N-terminal of huntingtin protein (HTT) for exporting HTT out of the nucleus. However, this interaction is significantly reduced by the polyQ expansion mutant HTT (mHTT), resulting in subsequent nuclear membrane distortions [76–78]. The identifications of NNMNAT2 interactomes located in the regulatory regions of EDEM3 and TPR, NMNAT2 associated genes (see Fig. 4 A-B for selected regions 1 and 2) and their significant correlated expressions with NMNAT2 in ROSMAP snRNAseq data set provide strong support for the validity of 4C-seq identified interactomes in regulating NMNAT2 transcription. The co-regulations of NMNAT2 and EDEM3 and TPR were further supported by our CRISPR-Cas9 ATF4, HSF1, and SOX11 knockdown experiments. We found that NMNAT2 and EDEM3 are co-regulated by HSF1 and SOX11 (Fig. 7 and Sup. Fig. 7-1). NMNAT2 and TPR are co-regulated by ATF4, HSF1, and SOX11. However, none of the tested TFs exert impacts on ODR4 mRNA levels.

Egl-9 family hypoxia-inducible factor (EGLN1) is crucial for iron homeostasis and its dysfunction is involved in the development of PD [79,80]. It has been reported that EGLN1 mRNA levels increase about 2-fold in the substantia nigra of PD brains [81]. Ring Finger Protein 2 (RNF2), encoding the enzymatic component of the PRC1 (polycomb repressive complex 1) complex, regulates the migration and differentiation of neural precursor cells and it is shown that its deletion triggers abnormal development of the central nervous system [82]. TRNA Methyltransferase 1-like (TRMT1L) is an RNA methyltransferase that is involved in RNA modification [83]. Dysfunction of RNA modification has been associated with neurological diseases [84–86]. TRMT1L is reported to regulate cognitive function, and its absence is shown to alter motor coordination and aberrant exploratory behavior in mice [87]. Pyrroline-5-carboxylate reductase 2 (PYCR2), encoding a mitochondrial enzyme, catalyzes proline biosynthesis by reducing pyrroline-5-carboxylate (P5C) into L-proline through NADH oxidation [88]. It is shown that depletion of PYCR2 leads to axonopathy and hypomyelination by excessively increasing cerebral glycine [89].

It remains to be determined why NMNAT2 and EDEM3/TPR expression are positively correlated in SH-SY5Y cells while negatively correlated in ROSMAP subjects. Future studies are required to demonstrate the co-regulations between NMNAT2 and its associated genes in healthy brains as well as to identify key upstream regulatory networks. It is also unclear whether the expressions of NMNAT2-associated genes are also altered in various conditions, such as brain injury or proteinopathy, when NMNAT2 mRNA levels are reduced.

### NMNAT2 TFs are related to cAMP signaling and cellular stress

A recent *in vivo* CRISPR-Cas9-based screening of 1,893 transcription factors using the mouse retina as a model system [8] identified ATF4 and SOX11 as critical transcription factors whose reduction promotes neuronal survival. Additionally, SOX11 overexpression resulted in the death of mouse α-retinal ganglion cells (α-RGCs) [90]. ATF4 has also been highlighted in the pathogenesis of AD [91,92]. Here we identified the presence of ATF4 binding motif on NMNAT2 promoter region and deleting either ATF4 or SOX11 increased NMNAT2 expression, indicating they can act as NMNAT2 transcriptional repressors. In glaucoma models, NMNAT2 overexpression increased the survival of retinal ganglion cells (RGC) [28]. Likewise, NMNAT2 overexpression attenuates tauopathy [93,69]. Given the neuroprotective role of NMNAT2 in axonal degenerative conditions, ATF4 and SOX11 may repress NMNAT2 expression and subsequently reduce neuronal survival.

The ATF/CREB family consists of a group of basic region leucine zipper (bZIP) transcription factors, which bind to the cyclic AMP response element (CRE) site. There are two main subtypes of CREB, including CREB/CREB1 and ATF4/CREB2 [94]. CREB functions as a transcriptional activator when it is phosphorylated and promotes memory-related synaptic plasticity [95]. Our previous studies showed CREB binding on the mouse NMNAT2 promoter [69] and identified several positive NMNAT2 modulators that are known to increase cAMP signaling [70]. Here we also found CREB/CREB1 as putative TFs (Supplementary Table 3) predicted from the 4C interactomes in undifferentiated and neuron-like SH-SY5Y cells. ATF4, on the other hand, suppresses synaptic strength and blocks long-term synaptic facilitation when over-expressed [96]. Deleting ATF4 increases synaptic strength and induces the growth of new synaptic connections [97]. In line with our previous finding that NMNAT2 abundance is positively correlated to the levels of synaptic proteins [21], synaptic deficits caused by ATF4 may be contributed by its suppression effect on NMNAT2. Moreover, it is found that ATF4 interacts with the CREB-binding protein (CBP) [98] and bZIP TFs often operate as obligate dimers to activate transcription ([99]. For example, ATF4 dimerizes with one of CREB3-like proteins, CREB3L2, to drive a transcriptional network of many genes differentially expressed in AD [100]. Whether the suppression of NMNAT2 is solely caused by ATF4 or by ATF4 interacting with CREB via CBP or a ATF4-CREB dimer, requires further study to elucidate.

HSF1 is a major transcription factor that induces the expression of heat shock proteins (HSP) for quality control of proteostasis in eukaryotes [101] and its activity is altered in Huntington’s disease [102]. Our present study revealed that HSF1 downregulation leads to up-regulated NMNAT2 expression, suggesting HSF1-mediated transcriptional regulation of NMNAT2. In Drosophila, HSF is the critical transcription factor for NMNAT gene expression under stress [103].

In summary, our study provides an unbiased identification of the transcriptional regulation of the human NMNAT2 gene. Identified genomic loci and TFs will provide us entrée points to further investigate how neurodegenerative insults reduce NMNAT2 mRNA levels and acquire a comprehensive view to slow neurodegenerative disease progression by upregulating NMNAT2 transcriptions.

## Supporting information

Supplementary Tables Description

Table1. Differentially expressed genes in undifferentiated and neuron-like SH-SY5Y cells

Table2. Transcription factor candidates from the NMNAT2 promoter

Table3. Transcription factor candidates from the interactomes of the NMNAT2 promoter

Table4. Motif occurrence (FIMO) analysis of the target transcription factors in region1

Table5. Motif occurrence (FIMO) analysis of the target transcription factors in region2

Table6. gRNA and primer sequences for piggyBac-based CRISPR/Cas9 gene deletion

## Data Availability

4C-seq and RNAseq data are deposited into NCBI-GEO (Gene Expression Omnibus) database.

## Acknowledgments

We thank the following: Dr. Xindan Wang and Dr. Cary Lai for helpful discussions; Dr. Marco Trizzino and Dr. J Mauro Calabrese for suggestions regarding the piggyBac-based CRISPR/Cas9 gene deletion method; the Center for Medical Genomics (CMG) School of Medicine Indiana University for 4C-sequencing service; Mr. Abhinav Bajpai and Mr. Scott Barton for technical assistance.

## Contributions

Y.C.C and H-C.L contributed to the conceptualization and experimental design. Y.C.C. conducted 4C-seq, the downstream bioinformatic analysis, piggyBac-Transposase modified stable cell line generation, knock-out/knockdown validation, and qPCR. S.Y. performed the TF knockdown cell line differentiation and qPCR. M.C. and K.N. analyzed the ROSMAP snRNA-seq dataset for gene co-expression. J-M.B. contributed intellectually to 4C-seq experimental design and downstream analysis. Y.C.C, S.Y., and H-C.L wrote the manuscript. All authors read, revised, and approved the final manuscript.

## Ethics declarations

The authors have no competing interests to declare that are relevant to the content of this article.

