## Supplementary Figures for ""Transcriptional Regulation of human NMNAT2: Insights from 3D Genome Sequencing and Bioinformatics""

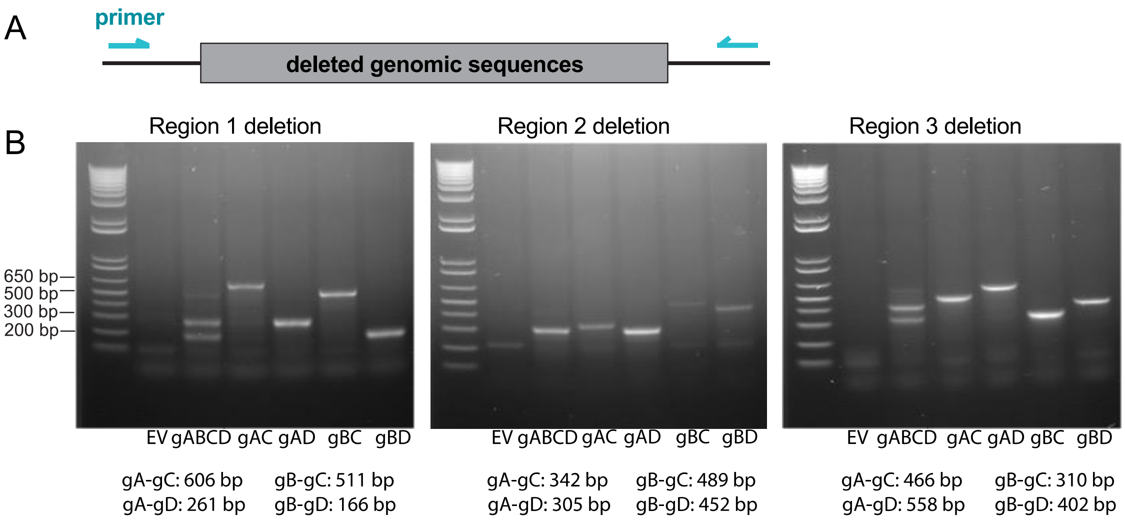


**Sup. Fig. 3-1. Genomic PCR to validate CRISPR-Cas9 deletions of the targeted genomic regions.** (**A**) PCR primers were designed flanking the deleted genomic sequences. (**B**) Representative DNA gel images show PCR products with sizes expected upon deletions. Abbreviations: EV, empty vector; gABCD, all the gRNAs; gAC, gRNA-A with gRNA-C; gAD, gRNA-A with gRNA-D; gBC, gRNA-B with gRNA-C; gBD, gRNA-B with gRNA-D. n = 3 independent batches of cells per group.

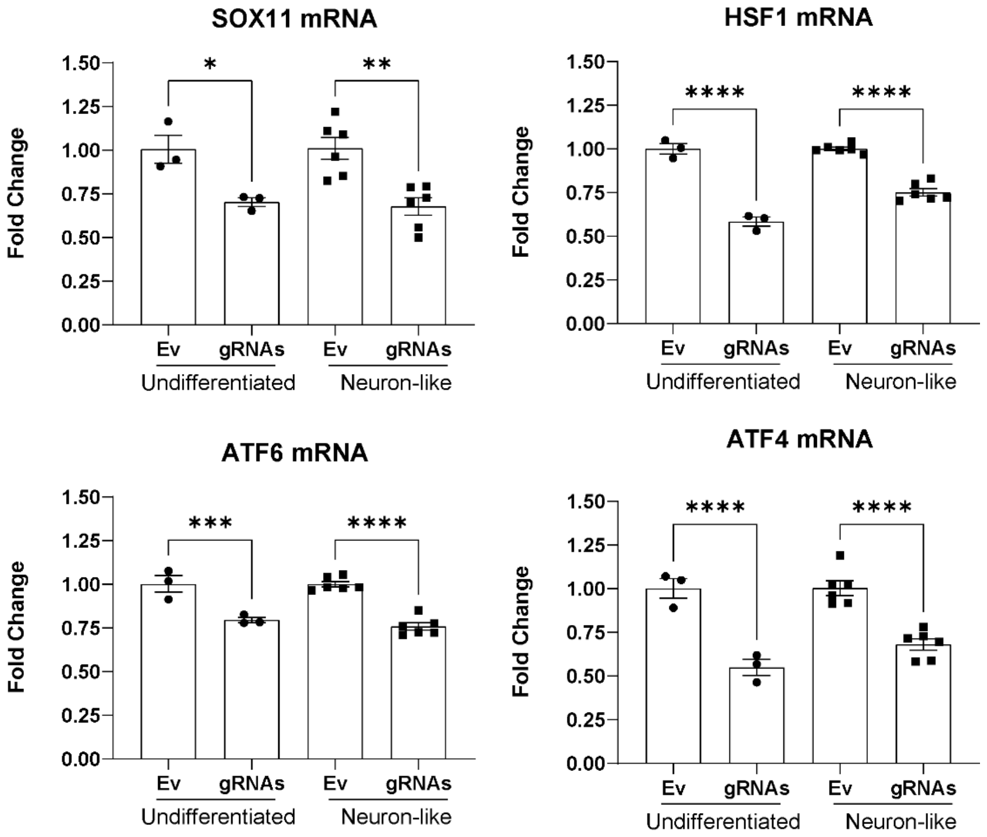


**Sup. Fig. 3-2. qRT-PCR to examine the knock-down efficiency with CRISPR-Cas9 deletion on SOX11, HSF1, ATF6 and ATF4.**

qRT-PCR was performed to determine the reduction of mRNA levels of the targeted TFs. Fold change was the ratio of mRNA levels after transfection with gRNAs compared empty vector (Ev) for both undifferentiated and neuron-like SY5Y cells. n=3 independent batches of cell culture per group. Data represents mean ± SEM. Ordinary one-way ANOVA with Šídák's multiple comparisons test was conducted, *p=0.0258, **p=0.0012, ***p=0.0004, **** p<0.0001.

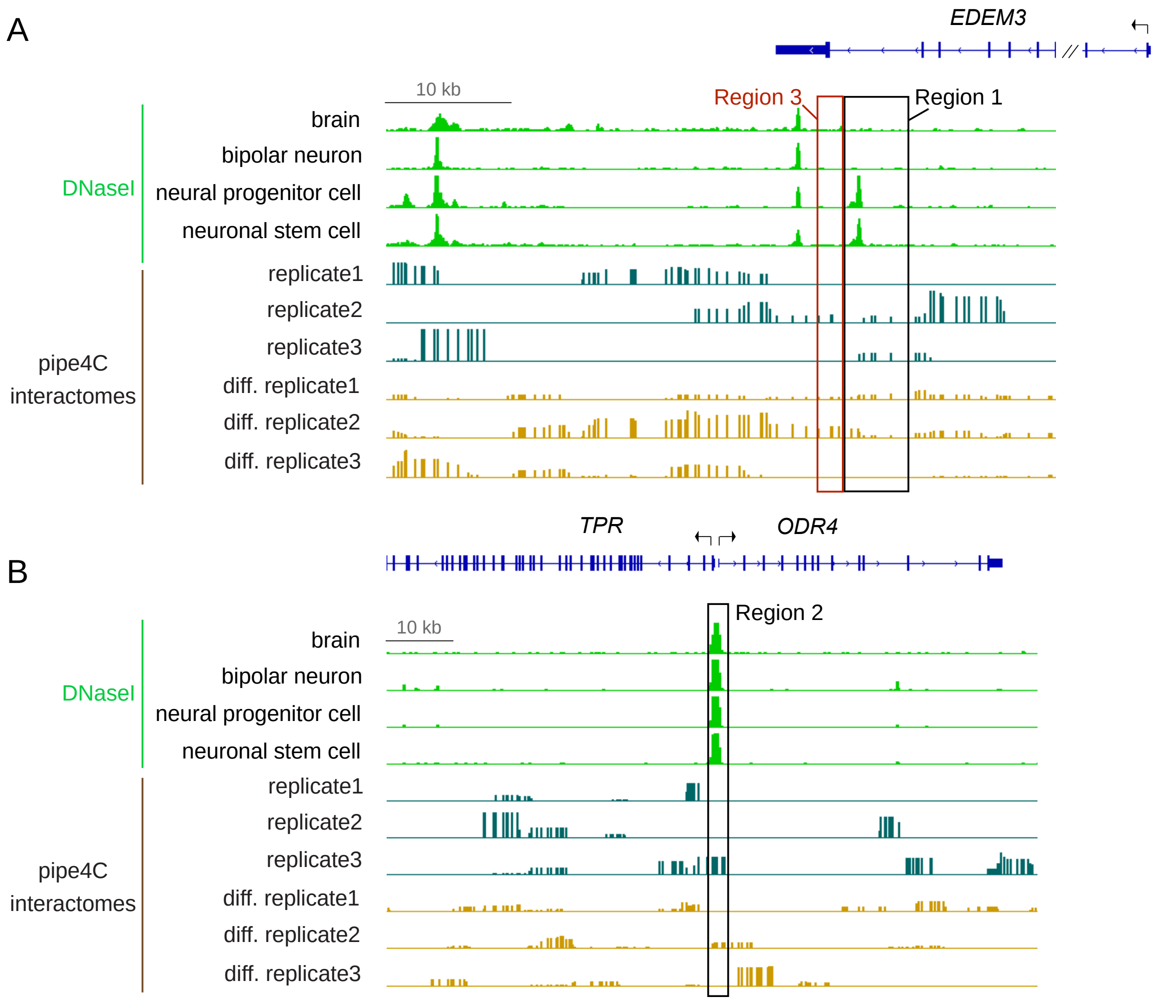


**Sup. Fig. 4-1. pipe4C interactomes in target regulatory regions.**
(A) pipe4C interactomes of three biological replicates in un-differentiated and neuron-like SHSY5Y cells aligned with the DNaseI peaks around the genomic regions of region1 and region3.
(A) pipe4C interactomes of three biological replicates in un-differentiated and neuron-like SHSY5Y cells aligned with the DNaseI peaks around the genomic region of region2.

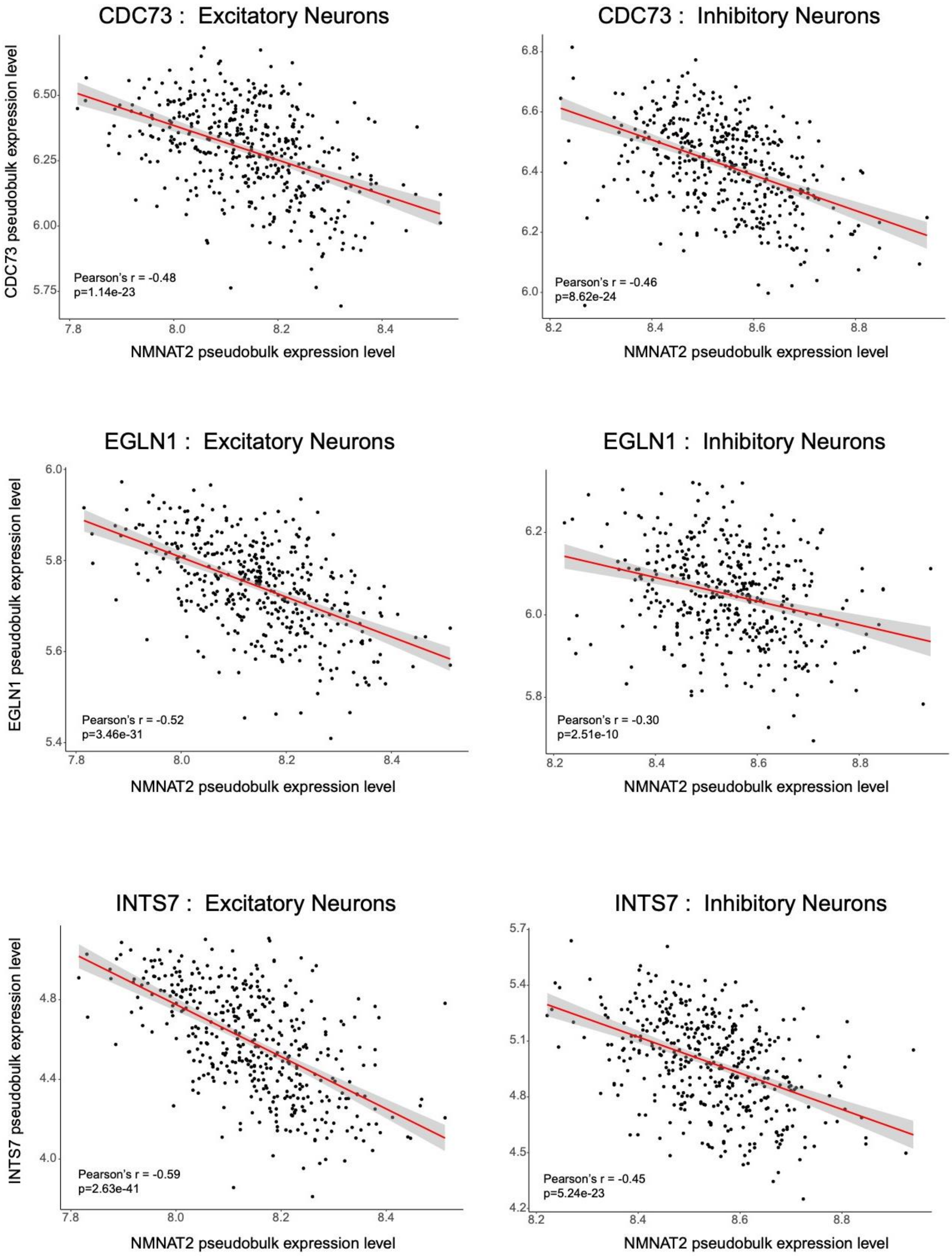
CDC73 : Excitatory Neurons
CDC73 pseudobulk expression level
Pearson's r = -0.48
p=1.14e-23
NMNAT2 pseudobulk expression level
CDC73 : Inhibitory Neurons
Pearson's r = -0.46
p=8.62e-24
NMNAT2 pseudobulk expression level
EGLN1 : Excitatory Neurons
EGLN1 pseudobulk expression level
Pearson's r = -0.52
p=3.46e-31
NMNAT2 pseudobulk expression level
EGLN1 : Inhibitory Neurons
Pearson's r = -0.30
p=2.51e-10
NMNAT2 pseudobulk expression level
INTS7 : Excitatory Neurons
INTS7 pseudobulk expression level
Pearson's r = -0.59
p=2.63e-41
NMNAT2 pseudobulk expression level
INTS7 : Inhibitory Neurons
Pearson's r = -0.45
p=5.24e-23
NMNAT2 pseudobulk expression level

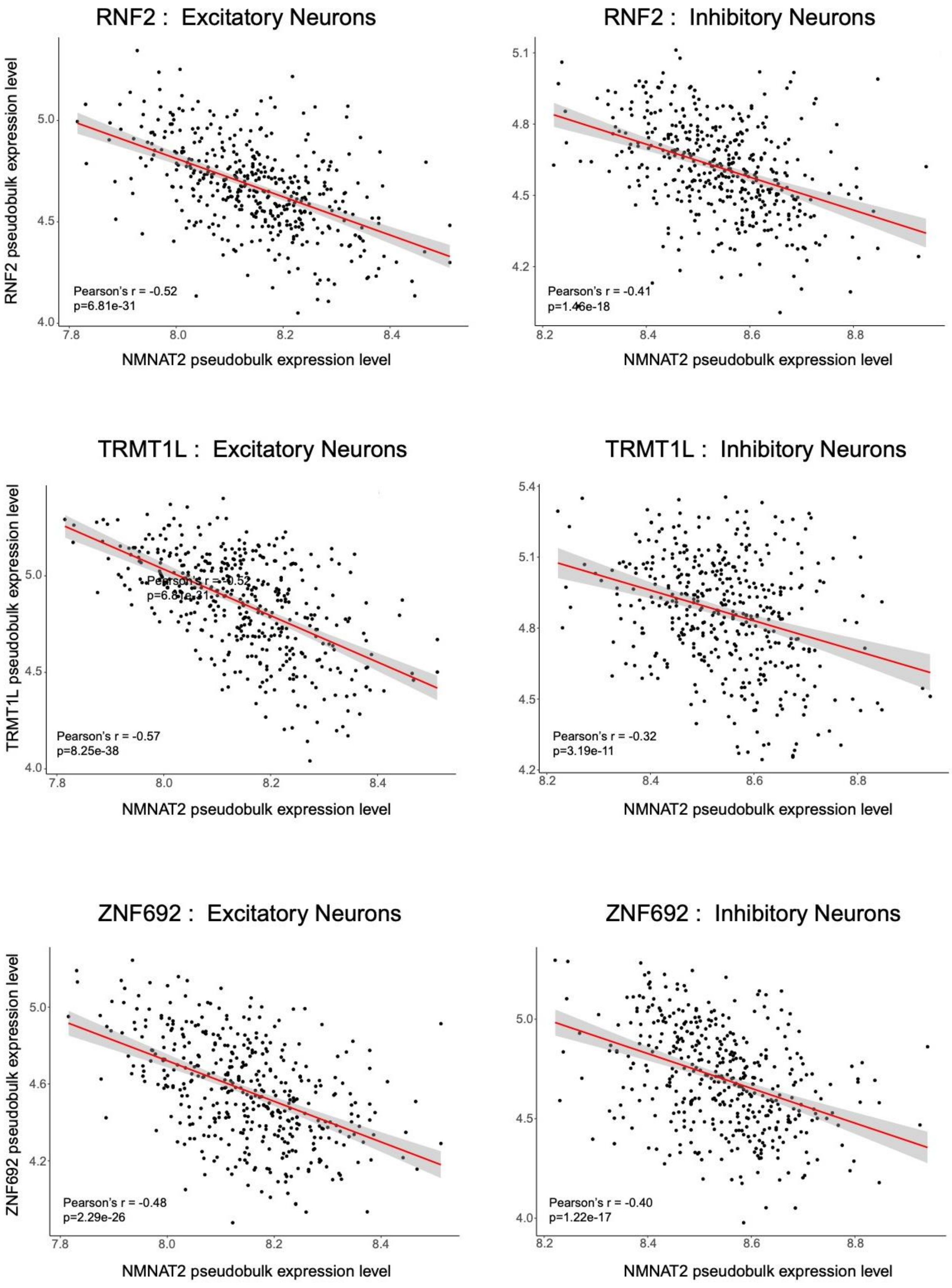

RNF2 : Excitatory Neurons
RNF2 pseudobulk expression level
Pearson's r = -0.52
p=6.81e-31
NMNAT2 pseudobulk expression level
RNF2 : Inhibitory Neurons
Pearson's r = -0.41
p=1.46e-18
NMNAT2 pseudobulk expression level
TRMT1L : Excitatory Neurons
TRMT1L pseudobulk expression level
Pearson's r = -0.57
p=8.25e-38
NMNAT2 pseudobulk expression level
TRMT1L : Inhibitory Neurons
Pearson's r = -0.32
p=3.19e-11
NMNAT2 pseudobulk expression level
ZNF692 : Excitatory Neurons
ZNF692 pseudobulk expression level
Pearson's r = -0.48
p=2.29e-26
NMNAT2 pseudobulk expression level
ZNF692 : Inhibitory Neurons
Pearson's r = -0.40
p=1.22e-17
NMNAT2 pseudobulk expression level

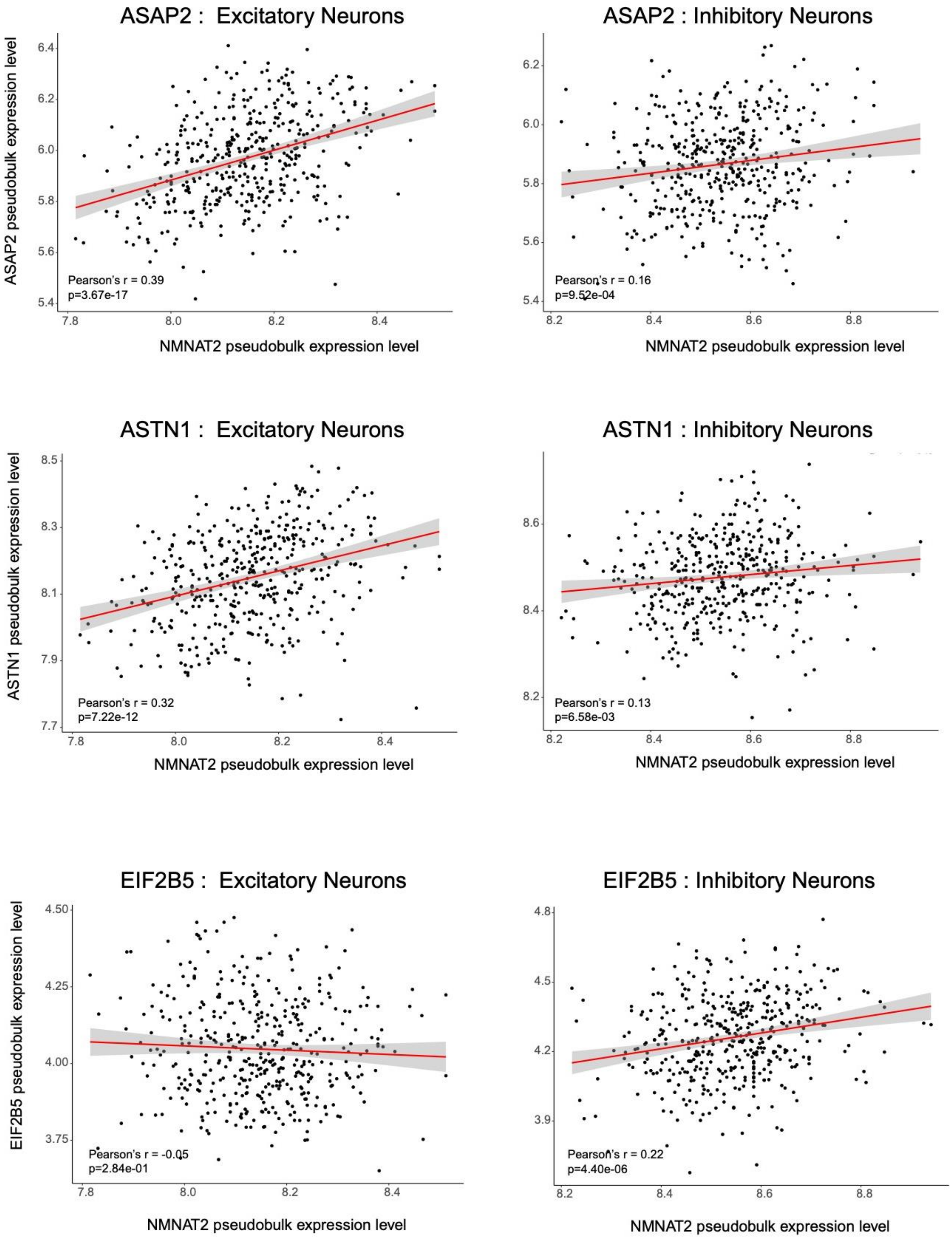

ASAP2 : Excitatory Neurons
ASAP2 pseudobulk expression level
Pearson's r = 0.39
p=3.67e-17
NMNAT2 pseudobulk expression level
ASAP2 : Inhibitory Neurons
Pearson's r = 0.16
p=9.52e-04
NMNAT2 pseudobulk expression level
ASTN1 : Excitatory Neurons
ASTN1 pseudobulk expression level
Pearson's r = 0.32
p=7.22e-12
NMNAT2 pseudobulk expression level
ASTN1 : Inhibitory Neurons
Pearson's r = 0.13
p=6.58e-03
NMNAT2 pseudobulk expression level
EIF2B5 : Excitatory Neurons
EIF2B5 pseudobulk expression level
Pearson's r = -0.05
p=2.84e-01
NMNAT2 pseudobulk expression level
EIF2B5 : Inhibitory Neurons
Pearson's r = 0.22
p=4.40e-06
NMNAT2 pseudobulk expression level

**Sup. Fig. 5-1. Relationships between NMNAT2 and the associated genes in snRNA data from the DLPFC of ROSMAP human subjects.**

Selective example of scatter plots showing the relationship between NMNAT2 mRNA levels to NMNAT2-associated genes: CDC73, EGLN1, INTS7, TRMT1L, ZNF692, ASAP2, ASTN1 and EIF2B5 in excitatory and inhibitory neurons. n = 424 human subjects.

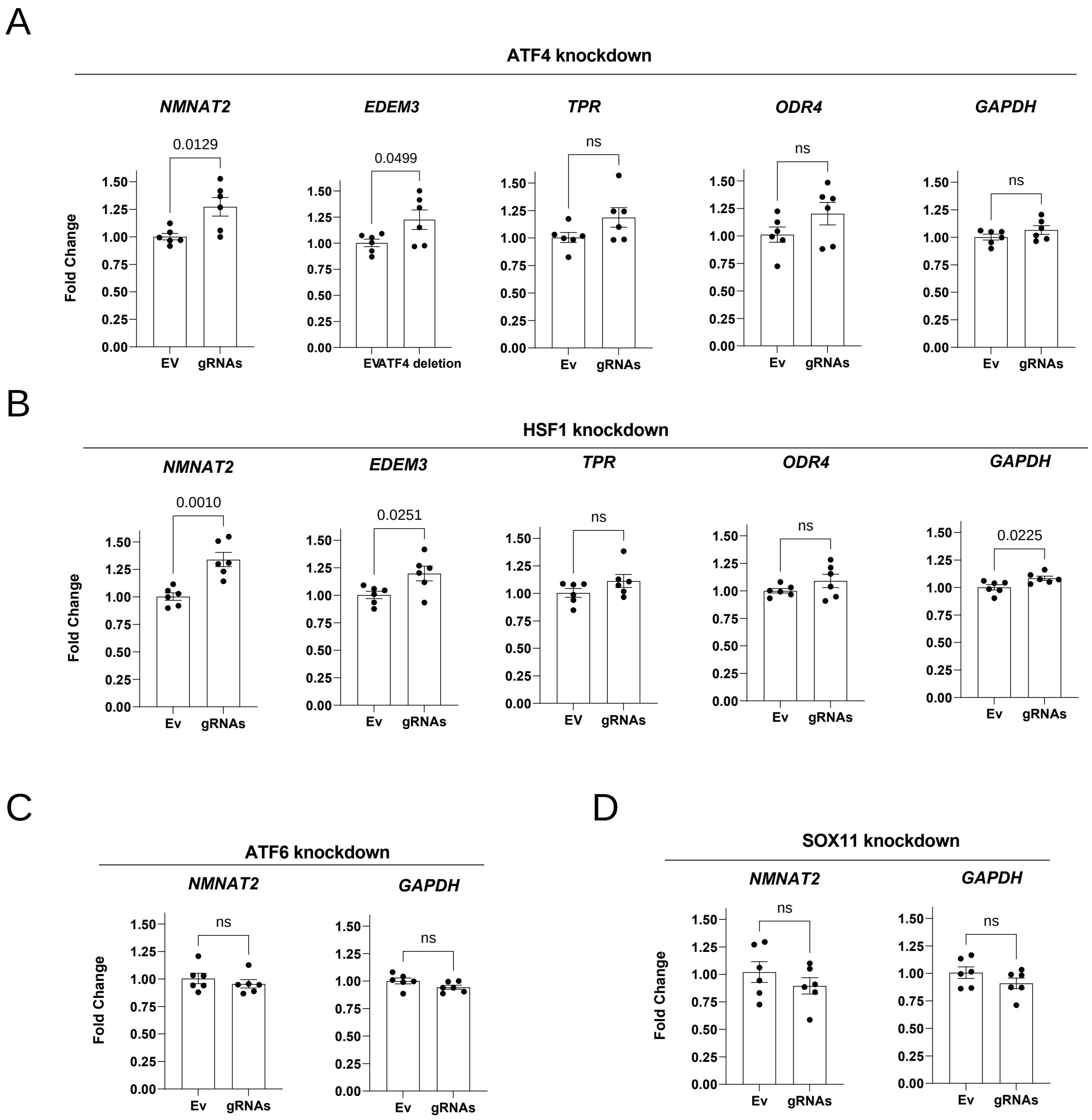


**Sup. Fig. 7-1. Gene expression of NMNAT2 and NMNAT2-associated genes upon knockdown of the target transcription factors in neuron-like SH-SY5Y cells.**

(A) ATF4 knockdown led to up-regulation of NMNAT2 and EDEM3 mRNA. (B) HSF1 knockdown led to up-regulation of NMNAT2 and EDEM3 mRNA. (C, D) ATF6 or SOX11 knockdown does not change NMNAT2 mRNA abundance. n = 3 biological replicates per batch of differentiation and 2 independent batches of differentiation per group. Data represents mean ± SEM. Unpaired student's t-test was conducted.
